# A comprehensive phage-bacteria interaction atlas links phage lineage and capsule serotype to genome-guided machine learning prediction in *Klebsiella pneumoniae*

**DOI:** 10.64898/2026.08.12.744533

**Authors:** Hemaa Selvakumar, Avery J. C. Noonan, Ella Rotman, Mohamad Alayouni, Denish Piya, Flavien Maucourt, Sarshad Koderi Valappil, Madeline Svab, Britney Oriheula, Gina Cowser, Isabella Murray, Collis Bousliman, Alexey Kazakov, Adam M. Deutschbauer, Simon Roux, Mark Mimee, Adam P. Arkin, Vivek K. Mutalik

## Abstract

*Klebsiella pneumoniae* is a WHO critical-priority pathogen for which strain-specific bacteriophages are being explored as precision antimicrobials, yet rapid phage-host matching remains a major barrier to therapeutic deployment. We constructed a comprehensive interaction atlas comprising 84 taxonomically diverse phages and 101 globally sourced, clinically representative *K. pneumoniae* strains, including multidrug-resistant isolates. Systematic pairwise profiling produced 8,484 interaction measurements, of which 2,656 (31.3%) scored positive for bacterial clearance. Genus was the dominant phage-side determinant of host range, while capsule K-serotype was the strongest host-side determinant of susceptibility; aggregate defense, prophage, plasmid, and antimicrobial-resistance features contributed comparatively little. A genome-guided machine learning model predicted interactions without curated host annotations (AUROC, 0.882; AUPR, 0.765), outperforming a model based only on phage genus and K-serotype and modestly exceeding a curated genomic baseline. The model recovered capsule- and lipopolysaccharide-biosynthesis genes, canonical receptors and defense-associated features as major predictors using SHAP analysis. Feasibility tests of expert- and model-selected cocktails exposed a translational constraint. Although all formulations suppressed growth *in vitro*, only the specific cocktail whose phages replicated robustly within the murine gut reduced colonization, suggesting *in vivo* amplification rather than predicted host range as the limiting factor for therapeutic efficacy. Together with the activity of a model-selected cocktail built for an isolate completely excluded from training, these results provide a species-wide resource for *K. pneumoniae* phage matching and support a hybrid workflow combining genome-based ranking with targeted phenotypic validation.

## Introduction

*Klebsiella pneumoniae* is a WHO critical-priority pathogen and a core member of the ESKAPE group, responsible for healthcare-associated pneumonia, urinary tract infections and bloodstream infections with high mortality^1–6^. A large and growing share of these infections is caused by strains that carry carbapenem and cephalosporin resistance together with hypervirulent genetic backgrounds, and for many of them conventional antibiotics no longer offer a reliable treatment option^3,4,7–11^. Bacteriophages have re-emerged as a way to treat these strains^8,12,13^.

Bacteriophages, viruses that infect and kill bacteria, have several properties that make them attractive therapeutic agents^14^. They exhibit stringent specificity at the level of the individual strain and act on the target pathogen without disrupting the surrounding microbiota; they replicate at the site of infection rather than being cleared like a fixed dose of drug; and many *Klebsiella* phages degrade the capsule and biofilm matrix that shield the cell^7,15–18^. Phages are also abundant and renewable, so new ones can be isolated as resistance emerges, and existing phages can be engineered or evolved to match a target^18–21^.

The outermost capsule of *K. pneumoniae* represents the primary barrier to infection and the main determinant of phage host range, which phages typically recognize and degrade using tail-fiber depolymerases^22–28^. Over 150 genetically distinct capsular types exist, and individual phages are usually restricted to a few specific types, meaning a phage infecting one strain of a given K-type will likely infect most others of that type^29–33^. Capsular type is therefore the strongest single predictor of susceptibility, though it remains insufficient on its own^29,34^. Strains sharing a K-type can carry different capsule biosynthesis alleles, causing variations in susceptibility, while infection also depends on sub-capsular receptors like lipopolysaccharide, the O-antigen and outer membrane proteins that specific phages target via non-depolymerase receptor-binding proteins^29,30,35,36^. Furthermore, post-adsorption defenses introduce an additional layer of resistance that capsular typing cannot capture. Consequently, predicting infection requires more than K-type identification, and resolving susceptibility reliably necessitates the systematic measurement of phage-host interactions across extensive panels of strains and phages^37–40^.

In practice, phages are still matched to strains one isolate at a time. Candidate phages are screened against each new clinical isolate by plaque and liquid assays, and because resistance in *K. pneumoniae* arises mainly through changes in surface receptors, effective preparations are typically cocktails of phages that engage independent receptors to limit cross-resistance.

Assembling those cocktails rests on the same per-isolate screening, so the characterization has to be repeated in full for every new isolate, and it does not scale to the timeframes that treatment requires. Predicting host range directly from clinical whole-genome sequences eliminates the need for prior phage propagation, receptor mapping or isolate-specific resistance profiling. Models trained on diverse datasets can infer susceptibility for new isolates, a framework recently validated for *Pseudomonas aeruginosa* and *Escherichia coli* through genomic prediction, cocktail design and in vivo testing^41,42^. In both species, surface receptor identity was the primary determinant of susceptibility, whereas defense systems modulated interactions at the specific phage-strain level^41,42^. Because genomic models do not transfer effectively across genera, a predictive resource for *K. pneumoniae* must be developed using *K. pneumoniae* data, rather than adapting models from other species^43^.

Progress in modeling *K. pneumoniae* host range is limited by dataset scale and architectural design constraints^44^. Historical interaction studies rely on small phage panels against reference strains or local isolates, failing to capture global species diversity and leading models to learn lineage-specific features rather than universal determinants^23,25,29,30,34,44–50^. Existing models also depend heavily on pre-selected features^43,44^. For instance, clustering depolymerases and receptor-binding proteins predicts capsule-mediated interactions but treats capsular type as a categorical label, failing to resolve strain-level variations in capsule biosynthesis loci or account for capsule-independent infection^29^. Recent approaches restrict inputs to specific capsule loci or employ protein-language-model embeddings of tail fibers. These methods maintain an exclusively receptor-centric perspective, remain vulnerable to severe data imbalance in sparse interaction matrices and utilize deep architectures with constrained interpretability^23,45^.

Here we constructed a complete 84-phage by 101-strain interaction atlas spanning globally sourced, drug-resistant clinical isolates^51^ and taxonomically diverse *Klebsiella* phages. We use the atlas to quantify host-range diversity, test network organization, partition interaction variation across host and phage features and evaluate our genome-guided machine learning prediction framework^41,43^. The results identify phage lineage and identity as major organizing axes, establish capsule serotype as the clearest host-side signal and show that genome-wide features add selective value for phenotypically atypical strains. We then used the model under distinct optimization specifications to design two phage cocktails and compared them with two expert-designed cocktails in liquid culture and a gnotobiotic mouse colonization model. Expert-selected cocktails delayed regrowth longest in culture, yet in mice only the specific cocktail whose phages amplified robustly within the gut reduced the *K. pneumoniae* burden. This outcome identifies *in vivo* amplification rather than predicted host range as the limiting variable for therapeutic success, while establishing the feasibility for genome-guided prioritization for unseen isolates. Ultimately, this work provides a practical framework that couples for genome-based phage selection with treatment-relevant validation against *K. pneumoniae*.

**Figure 1:**
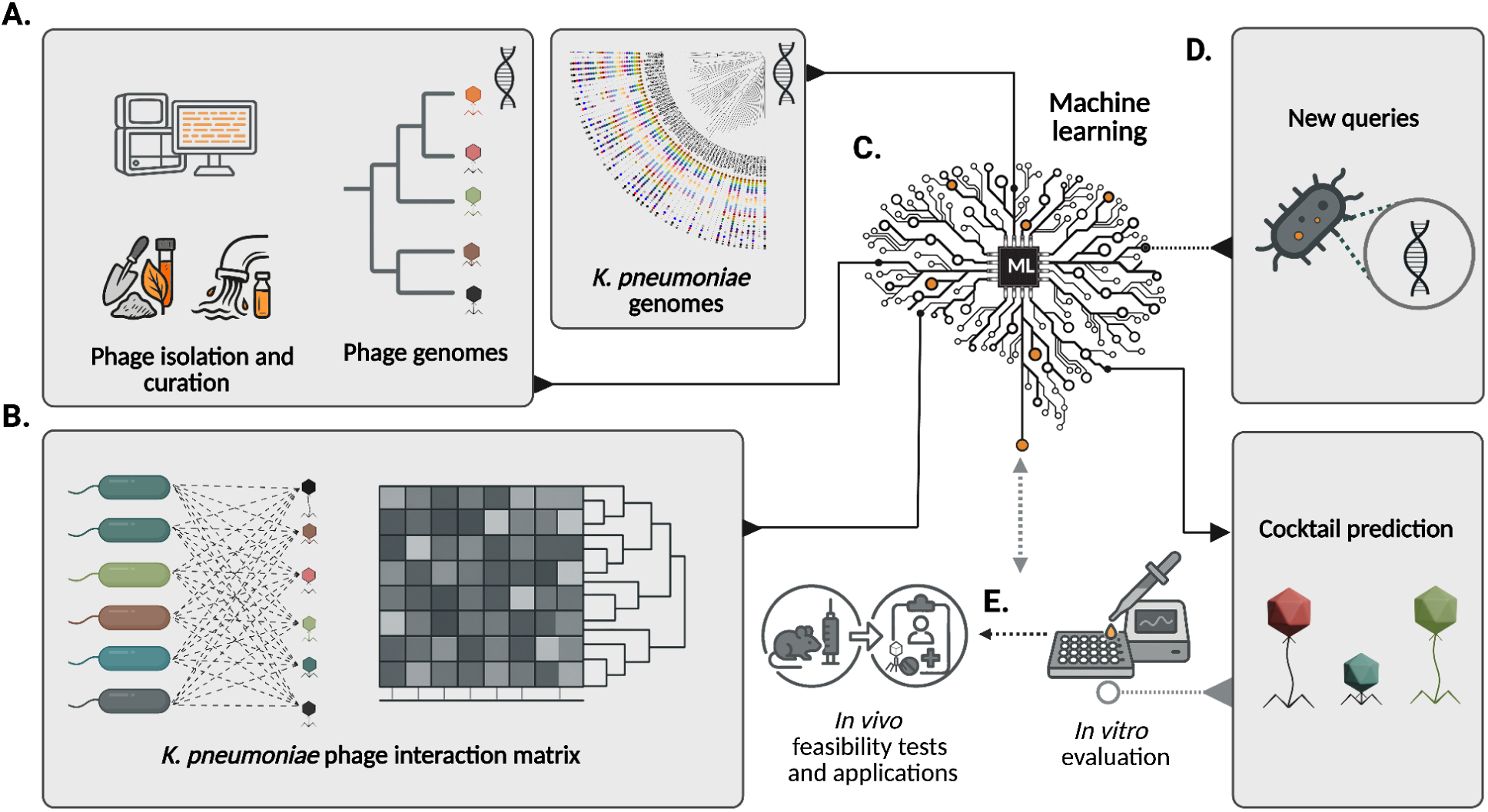
Overview of the ML-driven *K. pneumoniae* phage cocktail design pipeline. (A) Data collection and phage discovery: Isolation of bacteriophages from environmental sources, genomic sequencing and phylogenetic classification alongside strain genome sequencing. (B) Phenotypic interaction mapping: High-throughput screen generating a *K. pneumoniae*-phage interaction matrix and host range susceptibility profiles. (C) Predictive machine learning model: Integration of genomic and phenotypic datasets to train predictive algorithms for phage-host interactions. (D) Application and prediction: Querying uncharacterized clinical isolates to predict customized, synergistic phage cocktails. (E) Experimental validation: *in vitro* screen and *in vivo* animal model validation. Created in BioRender. X, A. (2026) https://BioRender.com/rgi8jma

## Results

### Assembly of a clinically diverse *K. pneumoniae* strain panel and a phage collection for mapping interactions

The bacterial panel comprised 100 well-characterized *K. pneumoniae* strains from the Multidrug-Resistant Organism Repository and Surveillance Network (MRSN) to ensure our interaction profiling would capture the breadth of genetic and phenotypic variation found in clinical populations^51^. This curated collection is a subset of 3,878 isolates collected between 2001 through 2020 from 63 healthcare facilities in 19 countries across multiple continents, offering exceptional geographic breadth and representation of global population structure (Supplementary figure S1). In this study, we have included an additional model strain MKP103, a derivative of strain KPNIH1, an early isolate from the NIH Clinical Center outbreak^52,53^. Together, this panel covers 94 sequence types, 54 K-loci, 11 O-antigen serotypes, and 74 of 101 strains were classified as multi-drug resistant or worse (Supplementary figure S2; Figure 2A). The genomic landscape of these isolates exhibited significant variation, carrying a median of 13 antimicrobial resistance genes, 24 phage defense and 6 anti-defense elements, and 5 prophages (Supplementary figure S3).

**Figure 2:**
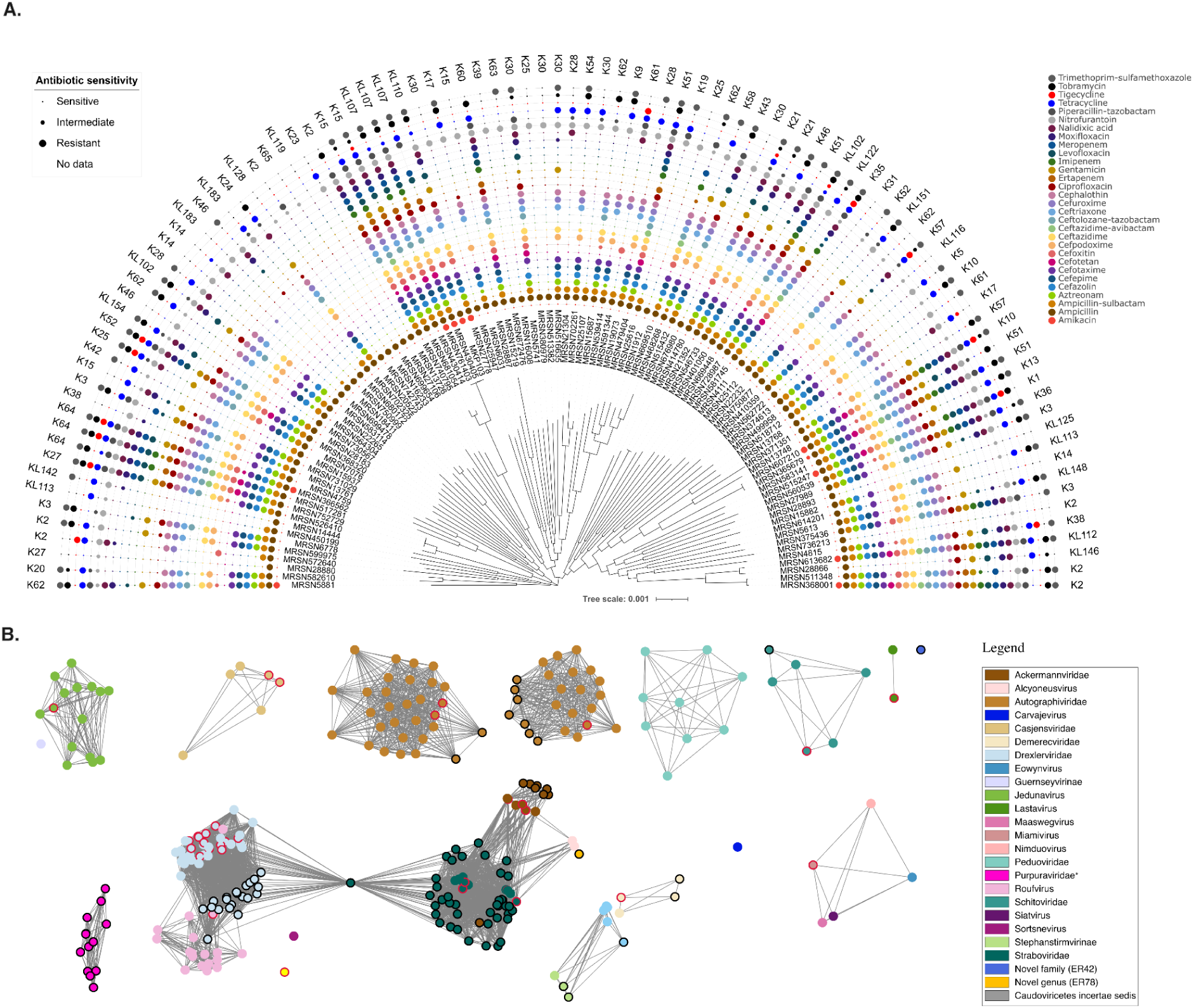
Genomic and taxonomic diversity of the *K. pneumoniae* strains and phages used in this study. (A) The neighbor-joining phylogenetic tree of 101 *K. pneumoniae* strains inferred from the multiple alignment of the core pangenome, showing the genomic diversity of the panel, with antibiotic resistance phenotype and isolation source annotated^51^; visualized in iTOL^56^. (B) Phage genome clustering network of 84 phages used in this study and 171 *K.pneumoniae* public phage genomes from the vConTACT reference database v230, generated with vConTACT3 and visualized in Cytoscape^57^. Phages from this study have black node borders and reference phages isolated or reported in the United States have dark red node borders; edges connect genome pairs sharing10 or more genes. Node colors and shapes denote taxonomic groups assigned by vConTACT3 as shown in the legend.

The phage panel comprises in-house isolates, phages from our previously described collection^54^ and the KlebPhaCol repository^35^. To maximize genomic and taxonomic diversity, we selected 84 phages and characterized their relationships using proteomic equivalence quotient (PEQ) scores based on the product of shared gene fraction and average amino acid identity (Methods, Supplementary figure S4). To contextualize our library within the global landscape of known *Klebsiella* phages, we generated a gene-sharing network comprising our 84 phages alongside 171 *K. pneumoniae* phage reference genomes from the vConTACT reference database v.230^55^ (Figure 2B). This panel spans 6 established viral families and 13 genera, with additional lineages lacking formal genus-level assignment, placing it among the most taxonomically diverse *Klebsiella* phage panels reported. For subsequent descriptions and analyses, annotation was standardized by assigning the family as the genus when the family was unassigned (specifically *Alcyoneusvirus* and *Marfavirus*) and by using the subfamily when present for unassigned genus (*Stephanstirmvirinae*).

### A complete interaction atlas captures broad phenotypic diversity

We characterized interaction outcomes across the complete matrix of 84 phages and 101 *K. pneumoniae* strains using systematic, high-throughput pairwise profiling (Methods).These assays capture bacteriostatic or bactericidal interactions that can arise from full lytic infection cycles, enzymatic or mechanical cell lysis, or growth inhibition. In this assay format clearance represents a pair-specific phenotype relevant to phage therapy applications. Interactions were scored on a three-point scale, where 0 is no visible clearance, 1 is hazy or turbid clearance indicating growth inhibition and 2 is a distinct zone indicating complete clearing, yielding 8,484 measurements.

Across the complete matrix, 2,656 pairs were positive (31.3%, Figure 3A and Supplementary Dataset 1). Per-strain interaction breadth or susceptibility was relatively evenly distributed (mean 26.3; median 25), so every strain was reached by multiple phages even though no single phage approached universal coverage. The per-phage interaction breadth or host range was markedly asymmetric and varied from 0 to 97 of 101 strains (mean 31.6; median 14), with two phages showing no detectable activity (Supplementary figure S5). Phages classified as *Marfavirus* and *Straboviridae* had the broadest host ranges, infecting a mean of 83 and 55 strains, respectively. In contrast, *Ackermannviridae* and *Autographiviridae* phages had narrow host ranges, infecting on average 4 and 7 strains, respectively. *Drexlerviridae* and *Purpuraviridae* occupied an intermediate spot. This specificity of the two non-interacting phage could stem from their initial isolation on *K. quasivariicola*.

**Figure 3:**
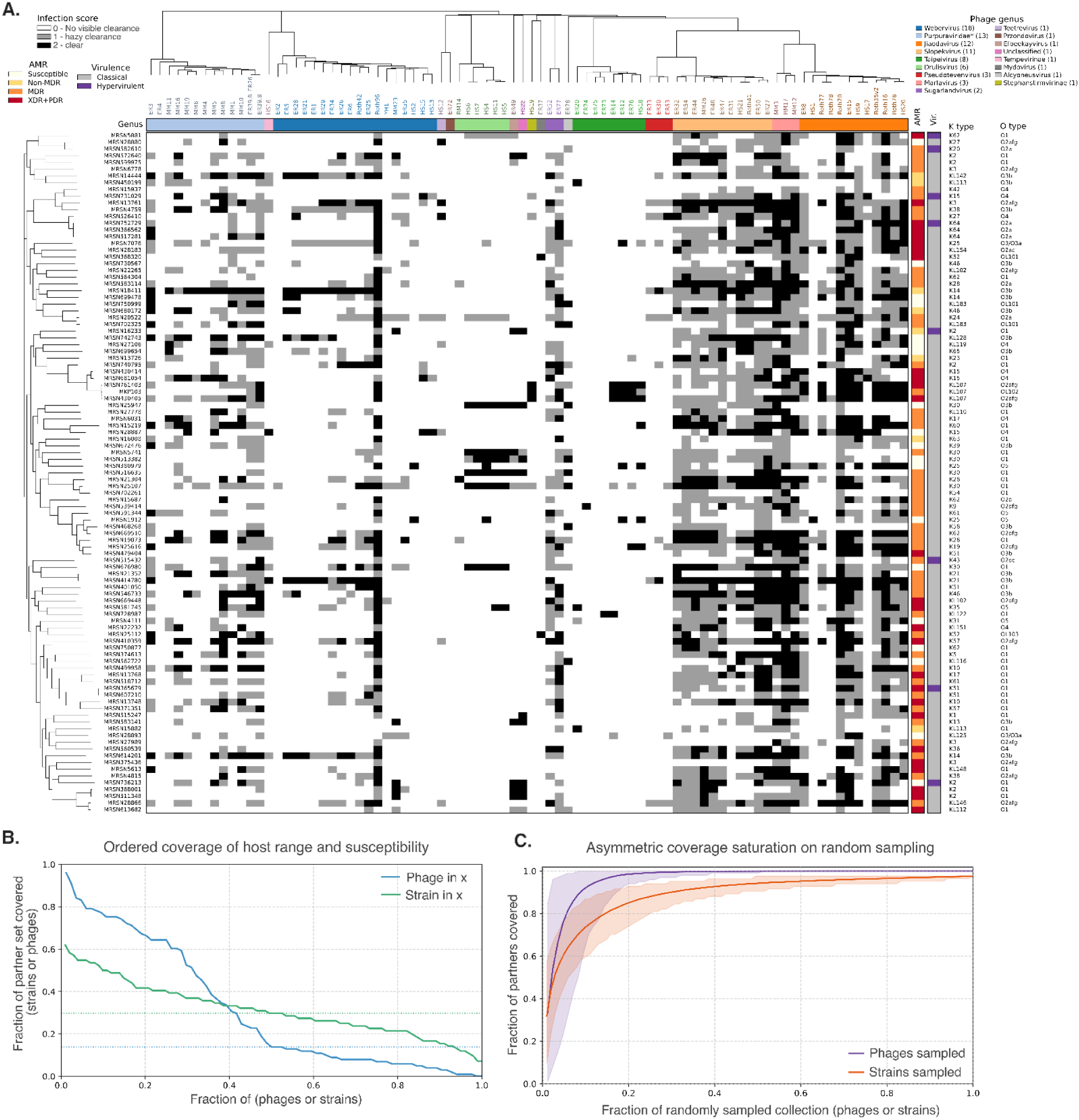
The complete *K. pneumoniae* phage-host interaction atlas and its coverage structure. A) Central heatmap of ordinal interaction outcomes for 101 *Klebsiella* strains against 84 phages. Rows are bacterial strains ordered by the strain phylogeny (left) and columns are phages ordered by the phage PEQ tree (above). B) Rank-ordered coverage curves for per-phage host range and per-strain susceptibility breadth. Fraction of strains cleared by each phage, ranked from broadest to narrowest; and fraction of phages clearing each strain, ranked from most to least susceptible. C) Rarefaction curves showing how interaction coverage accumulates when phages or strains are added in random order. The purple curve adds phages and measures the fraction of strains cleared by at least one phage; the orange curve adds strains and measures the fraction of phages with at least one detected host. Lines show the mean across 1,000 random permutations and shaded regions show the 95% permutation envelope.

Rarefaction analysis reinforced the distinction between broad coverage and complete coverage (Figure 3C). A small number of broadly infective phages covered most strains, but complete coverage required a substantially larger set containing complementary specialists. Conversely, a limited strain subset detected activity for most phages, while the two inactive phages remained unresolved in this assay panel. These distributions show that breadth is concentrated in a minority of phages and that a meaningful phage bank must retain narrow-range phages that reach otherwise poorly covered hosts.

### The *Klebsiella* phage-host interactions organize as permeable modules rather than a nested hierarchy

To evaluate the architectural hierarchy of the network, we performed a bipartite nestedness analysis on the interaction matrix, which exhibited a connectance of 0.32. This approach helped us to evaluate whether the hosts interacting with specialist phages formed proper subsets of those susceptible to generalist phages. The observed NODF metric was 64.85, compared with a degree-preserving null mean of 66.94 ± 0.29 (z = -7.30, Figure 4A, Methods). Thus, the apparent nestedness largely reflected differences in the numbers of hosts infected by each phage and phages infecting each host. After accounting for those degree distributions, specialist host ranges were not simply subsets of broader host ranges.

**Figure 4:**
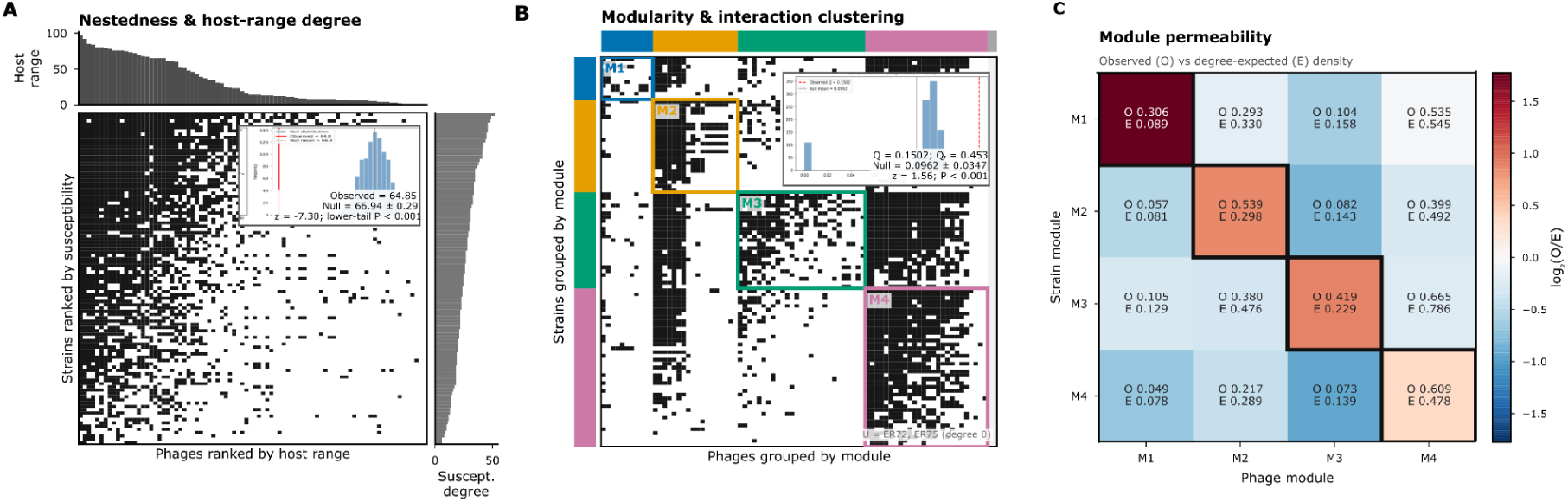
Organization and cross-module connectivity of the *K. pneumoniae*-phage interaction network. **(A)** Black cells show positive interactions between 101 strains and 84 phages, ordered from greater to lower per-strain and per-phage interaction breadth; nestedness was lower than expected after accounting for these breadth distributions (NODF, 64.85 versus 66.94+/- 0.29); P<0.001). **(B)** Reordering the same matrix revealed four LP-BRIM modules (Q=0.1502; P<0.001), with 45.3% of positive interactions occurring within corresponding strain and phage modules (Q_r_=0.453); two phages had no positive interactions and remained unassigned. **(C)** Each cell compares the observed interaction density (O) with that expected from the interaction breadths of the strains and phages (E); red indicates more and blue indicates fewer interactions than expected, while black outlines mark within-module comparisons.

To evaluate the higher-order organization of these interactions, we applied a bipartite modularity optimization framework via a complementary LP-BRIM analysis and resolved the network into four modules with Q = 0.1502, Qr=0.453 (Methods, Figure 4B, Supplementary figure S6). The observed modularity exceeded the degree-preserving null distribution (p < 0.001), showing that infections were concentrated within particular phage-strain groupings more strongly than expected from host range and per-strain interaction breadth alone. The modest value indicates that these groupings are permeable rather than strictly separated - cross-module infections occur, but less frequently than expected (Figure 4C). Together, these analyses indicate that phages and strains partition into loosely bounded interaction groups, suggesting that partially overlapping susceptibility and host-range determinants operate across this collection.

### Phage lineage and host capsule serotype organize interaction outcomes

To identify the key drivers of phage-host interaction phenotypes, we compiled a targeted set of phage- and strain-level features. Phage parameters comprised genus-level taxonomy, genome size, CDS count and tRNA gene content while bacterial traits included capsule and O-antigen serotypes, annotated defense and anti-defense repertoires, prophage load and plasmid-associated elements (Methods, Supplementary Datasets 2 and 3). These features were selected to capture core biological components of *K. pneumoniae* infection dynamics and they serve as an interpretable baseline to benchmark more complex, genome-based representations in downstream analyses.

Phage genus was the dominant determinant of host range (Kruskal–Wallis ε² = 0.6, Figure 5A) and remained significant after conditioning on proteome (PEQ) similarity (dbRDA, 24.8% unique variance). This suggests that while related phages tend to have similar host-targeting profiles, genus retains explanatory information beyond the proteomic similarity captured by the tree. Genome size, CDS and tRNA counts tracked host range univariately but not after controlling for genus, so they mark broad-host-range lineages rather than acting as independent predictors (Figure 5B, C). Explicit within-versus-between-genus regressions showed that nearly all observed variation in genome features occurred between genera but none of these traits predicted host range within genera (all p>0.431). Selecting one random representative phage from each genera confirmed that the positive between-genus direction was not driven by one particular representative. The pair-level models reinforce the dominance of phage-level structure. In GLMM, phage genus accounted for 31.0% of latent-scale variance in infection probability, while individual phage identity accounted for another 18.0%, giving a combined phage-side contribution of 49.0% (Methods, Figure 5G).

**Figure 5:**
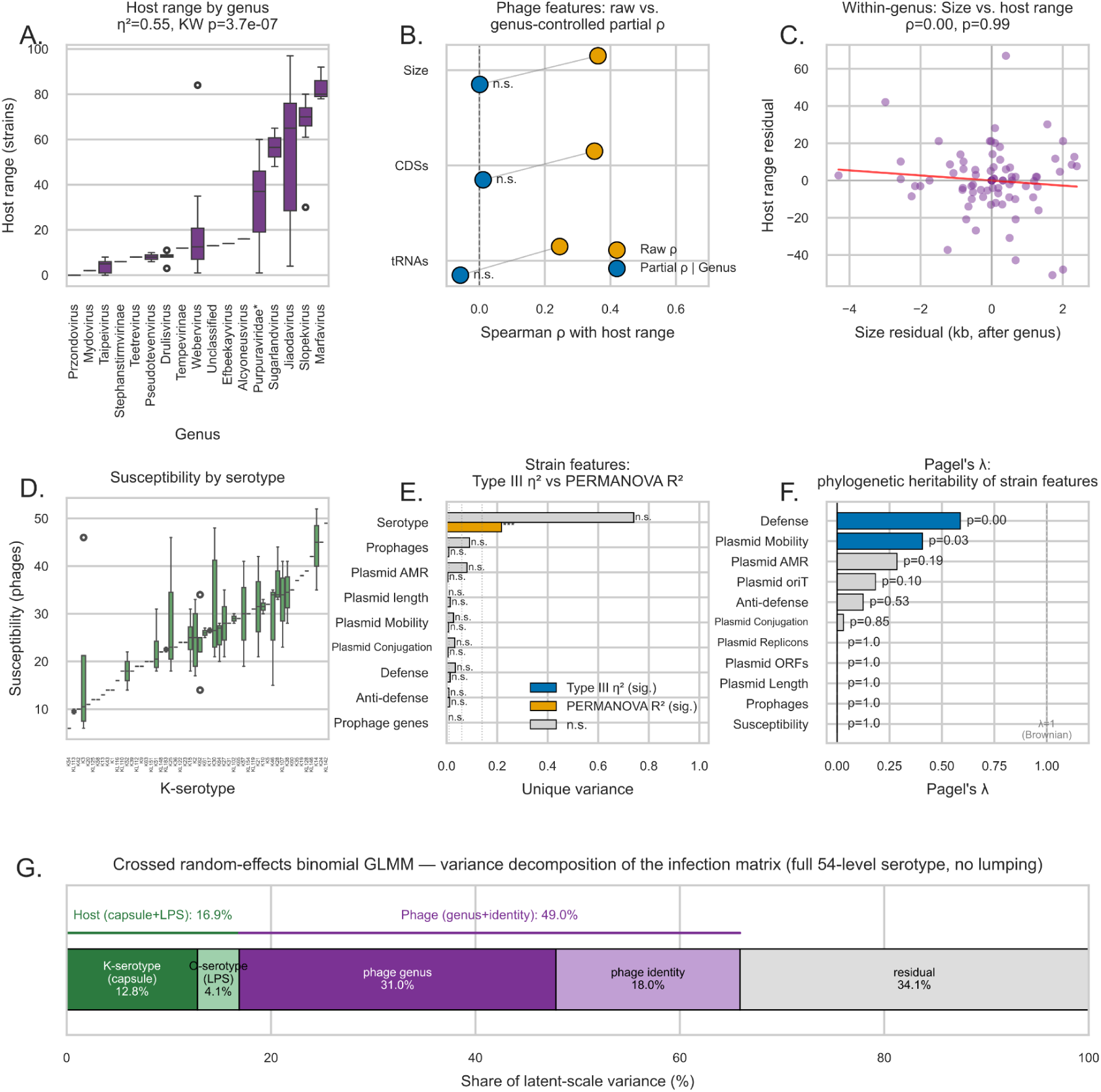
Phage and host determinants of *K. pneumoniae* infection across genus, serotype and other genomic features. (A) Phage host range variation by phage genus. (B) Phage genome size, CDS count and tRNA count were associated with host range in raw correlations, but these effects largely disappeared after controlling for genus. (C) Phage genus explained substantial variation in infectivity profiles, including after conditioning on PEQ similarity structure. (D) Strain susceptibility differed by K-serotype, with rare serotypes indicated separately according to the lumping rule used in the primary PERMANOVA/ANOVA analyses. (E) Variance metrics from strain-side multivariate models for K-serotype, defense, anti-defense, prophage and plasmid features. (F) Pagel’s lambda analyses showed little phylogenetic signal in overall susceptibility, but significant lineage structure in defense-system burden and plasmid mobility. (G) A crossed random-effects binomial GLMM fit to the full strain × phage infection matrix partitioned latent-scale variance across host and phage factors, showing that phage genus and individual phage identity dominated infection probability, while K-serotype explained more strain-side variance than O-serotype.

Host capsule (K) serotype was the dominant determinant of susceptibility (Figure 5D), explaining 19.1% of variation in the combined phylogeny-conditioned dbRDA (p=10^-4^). This result was consistent with the lumped-serotype susceptibility-count ANOVA (partial η²=0.269, p=0.024, Methods, Figure 5E) and with GLMM, in which K-serotype was the largest strain-side serotype component (12.8% of latent-scale variance; Figure 5G; Methods). O-serotype was a weaker and less robust secondary signal, it was significant in the lumped profile analysis but not in the susceptibility-count ANOVA or the full-resolution PERMANOVA sensitivity analysis, and is therefore treated as provisional. Each measured defense, prophage, plasmid or antimicrobial resistance (AMR) feature explained less than 5% of profile variation and was non-significant after accounting for the other predictors and phylogeny. Finally, Pagel’s λ found essentially no phylogenetic signal in the total susceptibility count (λ∼0, p=1). Therefore, closely related strains were not necessarily susceptible to similar numbers of phages. Some underlying features were phylogenetically structured, particularly defense count (λ=0.588, p=0.004) and plasmid mobility (λ=0.407, p=0.029), but their phylogenetic conservation did not translate into strong independent associations with susceptibility (Figure 5F). Collectively, these multiscale analyses demonstrate that phage-host infection outcomes are primarily governed by broad phage taxonomy and bacterial surface serotypes, whereas intracellular defense systems and host ancestral phylogeny exert minimal independent influence on overall susceptibility profiles.

### Genome-based machine learning robustly predicts interactions and recovers capsule signals without prior annotation

Identity and annotation analyses indicated that pairwise modeling was needed to resolve phage-host interactions. A two-way analysis attributed 39.85% of interaction variance to phage identity and 7.31% to strain identity (combined R² = 0.47, Methods), leaving more than half pair-specific or assay-associated. Likewise, multivariate tests associated serotype and phage genus with interaction phenotypes, but these features explained only part of the variance. We therefore tested whether models capturing nonlinear interactions could improve prediction and identify additional genetic correlates.

We compared three classifiers for predicting the susceptibility of an unseen *K. pneumoniae* strain to a fixed phage bank. Two reshuffled 10-fold cross-validation runs held out 10% of strains per fold, with feature selection, hyperparameter tuning, and fitting restricted to training data. Genus + K-Serotype included only phage genus and capsule serotype. The curated baseline added defense- and anti-defense-system counts, prophage burden, plasmid load, AMR phenotype, and genome-scale features. GenoPHI replaced these annotations with model-selected MMseqs2 pangenome protein clusters.

Across pooled held-out predictions (Figure 6A,B), Genus + K-Serotype performed worst (AUROC 0.810; AUPR 0.616), whereas the curated baseline (0.869; 0.735) and GenoPHI (0.882; 0.765) were closely matched. Thus, GenoPHI predicted strain-level interactions from genome-derived features without curated host annotations. Adding O-antigen serotype to the minimal model had little effect (AUROC 0.813 versus 0.810), consistent with capsule being the principal strain-side determinant.

**Fig. 6.**
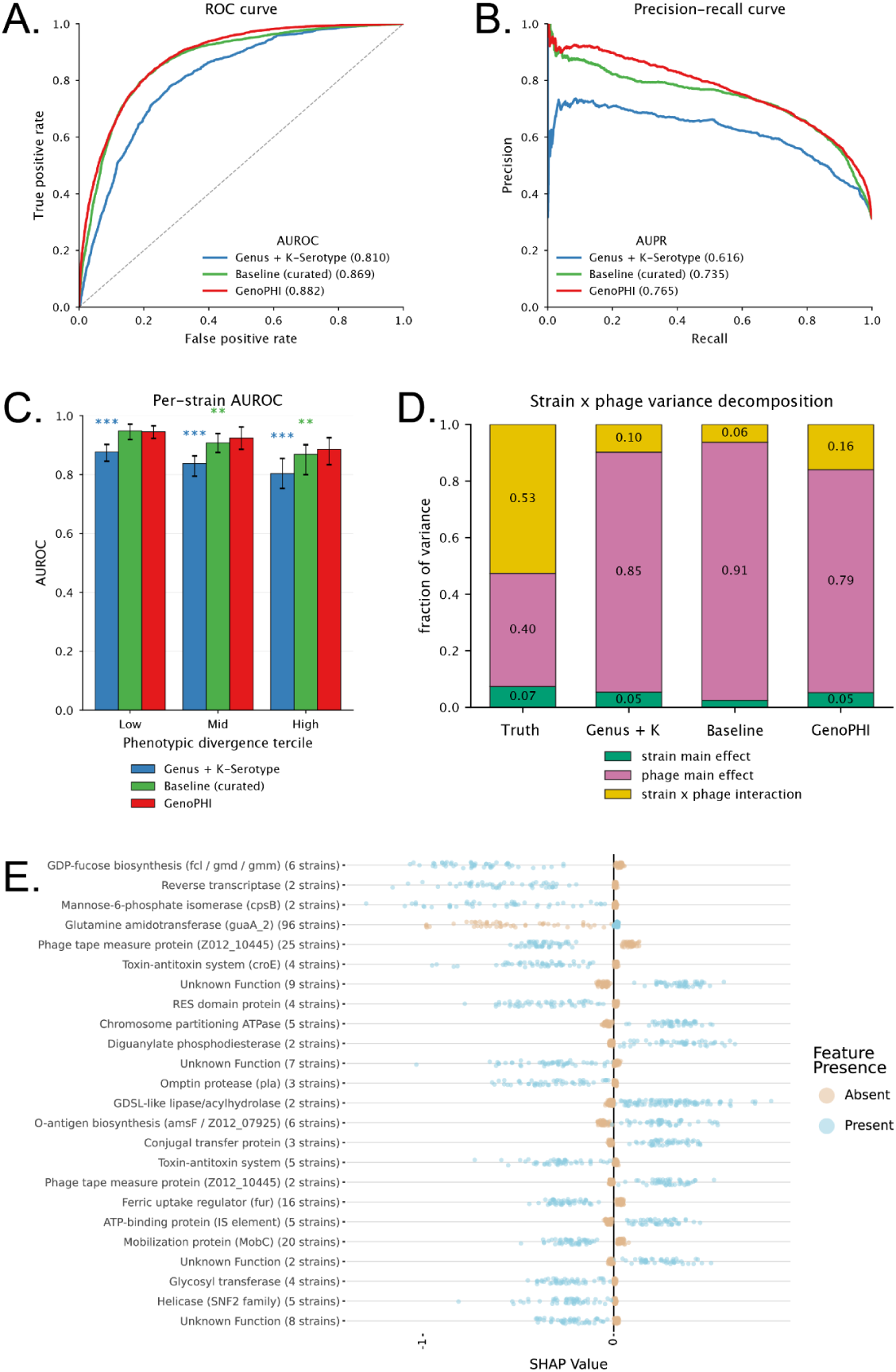
Machine learning-based strain-level *K. pneumoniae* phage-host interaction prediction. Three deployable models are compared throughout: Genus + K-Serotype (blue), curated baseline (green), and GenoPHI (red), evaluated under strain-held-out cross-validation (10 folds, two independent reshufflings; 20 fold-fits total). (A) ROC and (B) precision–recall curves on predictions pooled across all 20 held-out folds; legend entries give pooled AUROC and AUPR. (C) Per-strain held-out AUROC stratified by phenotypic divergence tercile. Bars show the median across strains; error bars span the interquartile range. Asterisks denote Holm-corrected paired Wilcoxon significance versus GenoPHI within each tercile (* p < 0.05, ** p < 0.01, *** p < 0.001). (D) Two-way (strain × phage) variance decomposition of the true interaction matrix and of each model’s held-out predictions, partitioning score variance into strain main effect, phage main effect, and the strain × phage interaction term. (E) SHAP beeswarm of the 25 highest-ranked strain-side feature clusters for GenoPHI, annotated with eggNOG-mapper gene-function calls. Each point is one modeling run’s median SHAP value for that cluster (50 runs), shown separately for the present (blue) and absent (orange) states. Positive SHAP values indicate the feature increases predicted infection probability; negative values indicate decreased probability. Strain counts in parentheses give the number of strains in which each cluster is present.

Entity-level analyses revealed different patterns (Supplementary figure S7). When ranking phages for a held-out strain, GenoPHI modestly improved median AUROC over the curated baseline (0.924 versus 0.912; Holm-corrected paired Wilcoxon p = 0.025), but not AUPR (0.859 versus 0.823; p = 0.07); both outperformed Genus + K-Serotype (AUROC 0.845; AUPR 0.648; both p ≤ 8 × 10⁻¹⁶). When ranking held-out strains for each phage, GenoPHI more clearly outperformed the curated baseline (AUROC 0.672 versus 0.587; p = 3 × 10⁻⁶) and the minimal model (0.571; p = 3 × 10⁻⁶). Because phages were not held out, this measures host-range resolution for known phages, not generalization to novel phages.

GenoPHI’s advantage concentrated among harder strains. Difficulty was measured by two largely independent axes (Spearman ρ = 0.04): phylogenetic novelty and phenotypic divergence from the five nearest phylogenetic training neighbors. The advantage was strongest and most orderly with phenotypic divergence (Figure 6C). GenoPHI and the curated baseline were indistinguishable among low-divergence strains (AUROC 0.945 versus 0.948, p = 0.34; AUPR 0.924 versus 0.927, p = 0.29), except for a small but significant curated-baseline MCC advantage (p = 3×10⁻⁴). GenoPHI’s AUROC advantage emerged at middle divergence (0.924 versus 0.908; p = 1.7 × 10⁻³), persisted at high divergence (0.886 versus 0.869; p = 1.2 × 10⁻³), and increased continuously with divergence (ρ = 0.24; p = 0.018). The novelty trend was weaker because all held-out strains had close training relatives (maximum nearest-neighbor distance 7.6 × 10⁻³ versus median pairwise 1.0 × 10⁻²; Methods). Variance decomposition supported this pattern (Figure 6D): pair-specific interactions comprised 52.6% of observed variance, and GenoPHI recovered 32.4% of this signal versus 17.2% for the curated baseline, while both assigned most remaining variance to the phage main effect.

SHAP analysis showed that GenoPHI recovered biologically coherent signals without serotype or receptor-location priors (Figure 6E). Predictive features included capsule and lipopolysaccharide biosynthesis genes (*fcl*, *gmd*, *gmm*, *cpsB*, and *amsF*), a glycosyltransferase, and canonical receptors (*fhuA* and *lamB*). Defense- and prophage-related features included a CroE-family antitoxin, a RES-domain protein, and phage-derived tape-measure proteins. GenoPHI also identified features absent from the curated representation, including a GDSL-like lipase/acylhydrolase, the omptin protease *pla,* and a diguanylate phosphodiesterase. Although their roles remain unknown, these genes are candidates for GenoPHI’s additional resolution on phenotypically divergent strains (Supplementary figure S9).

### Phage cocktail design for applications: Comparison of empirical selection and machine learning approaches

Phage-cocktail design for high-priority pathogens like *K. pneumoniae* is often constrained by incomplete knowledge of host receptors, environmental stability, and immune clearance. To evaluate cocktail selection strategies on a clinically representative yet biosafe host, we targeted MKP103, a carbapenem-sensitive derivative of the globally dominant ST258 clinical lineage (KPNIH1). We compared four distinct three-phage cocktails targeting MKP103, comprising two empirical-selection (ES1 and ES2) candidates chosen by domain experts and two variants selected using genome-guided machine learning (ML1 and ML2).

For empirical selection, eight phages demonstrating the strongest measured interactions with MKP103 were screened in liquid culture. Taxonomy and host range subsequently served as proxies for functional diversity due to limited receptor assignments. The ES1 cocktail comprised ER15 (*Jiaodavirus*), HS19 (*Stephanstirmvirinae*), and ER77 (*Sugarlandvirus*), whereas ES2 comprised ER15, HS18 (*Taipeivirus*), and ER39.8 (*Purpuraviridae*). Genetic screening, deletion, and complementation in ST258 identified LPS as the receptor class for ER15 and OmpC as the receptor for ER39.8^54^. The activity of ER39.8 was also modulated by LPS or O-antigen architecture, and with adaptive tail mutations restoring adsorption to resistant envelope mutants. Taxonomic precedent suggests that HS19 recognizes capsular polysaccharide, ER77 uses a capsule-dependent carbohydrate-recognition pathway, and HS18 recognizes a K-type capsule through tail-associated binding proteins or depolymerases^58–60^. However, these taxon-informed expectations are not phage-specific receptor assignments.

The ML1 cocktail comprised Roth20 and Roth16 (both *Jiaodavirus*) alongside RM17 (*Marfavirus*), while ML2 comprised Roth41 (*Slopekvirus*), ER39.8, and RM17. These formulations were selected using the cocktail-design procedure from GenoPHI with the specific target strain MKP103 completely excluded from the model training. The ML1 variant was generated via the standard GenoPHI workflow, which utilizes unsupervised HDBSCAN clustering to group phages by their predictive feature content before selecting top-performing candidates across distinct clusters. Conversely ML2 was derived from hierarchical clustering to partition phages into three predictive-feature groups to enforce broader taxonomic representation than HDBSCAN-based clustering. This was followed by liquid-culture screening of the three highest-probability candidates per cluster, which were ultimately narrowed to three phages based on individual performance. This secondary workflow leverages model predictions to identify candidate phages for direct validation against the target strain. Both ML1 design and ML2 candidate selection operate independent of prior interaction data for MKP103, rendering them immediately deployable to newly sequenced bacterial isolates.

### *In vitro* evaluation

All cocktails were evaluated in overnight liquid cultures against wild-type MKP103 and the capsule-deficient escape mutant ER649 (MKP103 *wzc*::IS5). The ES1 formulation demonstrated the highest efficacy, maintaining OD600 near baseline for 16 h without detectable bacterial regrowth. In contrast, ES2, ML1, and ML2 delayed MKP103 growth for only 8.5 to 11 h, resulting in AUCs of 2.4-3.4 OD·h and endpoint ODs of 0.5. Against ER649 mutant, ES2 provided moderate suppression (AUC 3OD·h; endpoint OD, ∼0.23), whereas ML1 and ML2 permitted rapid regrowth (endpoint OD, ∼0.8). The ES1 cocktail also consistently outperformed the individual phages in it. Conversely, ML2 provided minimal additional benefit over ER39.8 alone in WT MKP103 and exhibited lower efficacy than either ER39.8 or Roth41 against ER649. These findings indicate that expert-informed cocktails, particularly ES1, produced consistent suppression during *in vitro* testing.

### *In vivo* feasibility test

The cocktails were subsequently evaluated in gnotobiotic Swiss Webster mice colonized with a defined 13-species community^54^ (Methods). Mice received ∼5 x 10^7^ MKP103 CFU by oral gavage on Day 9, resulting in a median fecal colonization burden of 5 x 10^8^ CFU/g after four days. On days 14 and 15, separate groups of 3 to 5 mice were housed by treatment arm and administered 5% sodium bicarbonate followed by either PBS or a specific cocktail formulation (100 μl; 1 × 10^10^ PFU/ml). Fecal outputs were monitored on days 15, 16, and 18 with cecal contents harvested on day 21. Only the ML2 cocktail significantly lowered fecal MKP103 burden, achieving a 0.5 log₁₀ CFU/g reduction on day 15, which was maintained through days 16 and 18.

This therapeutic efficacy closely tracked fecal phage recovery, as ML2 alone exceeded 2 × 10^8^ PFU/g. The remaining cocktails yielded lower recoveries ranging from 10^4^ to 10^6^ PFU/g, despite an equivalent back-titrated delivery input of 10^10^ PFU/ml. These low recovery values point toward limited early gut amplification rather than delivery failure, though they may also be influenced by cage-specific environmental variations. Because both ES2 and ML2 contained identical quantities of ER39.8, a phage previously shown to maintain high stability in the murine intestinal tract^54^, its presence alone does not explain the distinct *in vivo* amplification profiles observed between the formulations. Although the underlying mechanisms preventing the other three cocktails from replicating robustly *in vivo* remain unresolved, especially given their successful host inhibition *in vitro*, these results demonstrate that an automated cocktail assembled without prior training data on a specific isolate can successfully achieve a modest reduction of a target pathogen within a complex colonized gut environment.

**Figure 7:**
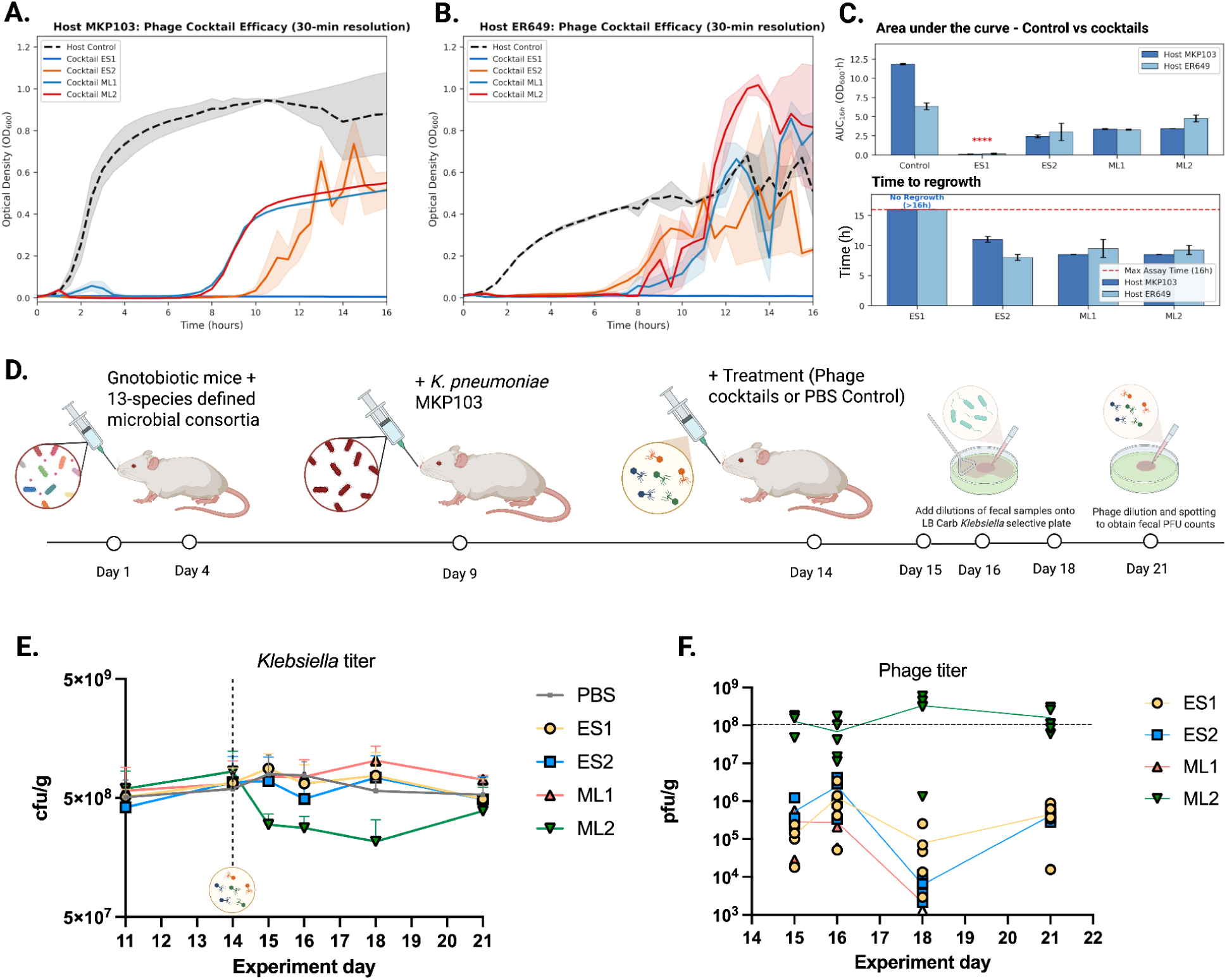
I*n vitro* killing kinetics and *in vivo* efficacy of phage cocktails against *K. pneumoniae*. (A,B) Bacterial growth curves of host strains MKP103 (A) and ER649 (B) under untreated control conditions (dashed black line) or challenged with ES or ML phage cocktails. (C) Top: Area under the curve (AUC) showing significant growth inhibition across treatments compared to control (p < 0.0001). Bottom: Time to bacterial regrowth within the 16-hour assay window; cocktail ES1 achieved complete growth suppression without detectable regrowth (>16h, horizontal dashed line). (D) Experimental timeline for the murine gut colonization model, illustrating administration of a defined microbial consortia, colonization with MKP103, and treatment with phage cocktails or PBS control. (E) Bacterial burden (cfu/g) measured in fecal samples prior to and post treatment (PBS control and phage cocktails), indicated by vertical dashed line. (F) Temporal dynamics of fecal phage shedding (pfu/g) across treatment groups (ES1, ES2, ML1, ML2) quantified on Days 15, 16, 18 and 21 following treatment. Horizontal dashed line denotes the minimum threshold associated with an observable decrease in fecal bacterial burden based on prior work^54^. Created in BioRender. X, A. (2026) https://BioRender.com/rgi8jma.

## Discussion

Phage therapy offers meaningful clinical promise for *K. pneumoniae*, particularly as a personalized adjunct for refractory multidrug- or carbapenem-resistant infections, though it remains an experimental strategy rather than standardized clinical care. A central obstacle is the strain dependence of phage activity, which requires therapeutic phages to be matched precisely to an individual patient’s infecting isolate. To evaluate whether bacterial genome sequences can guide this selection process, we built an experimental dataset comprising 8,484 interactions between 101 clinically representative strains and phages spanning 17 taxonomic lineages. The resulting network indicates that susceptibility is structured principally by phage lineage and bacterial surface architecture, while genome-wide features resolve critical phenotypic variations missed by aggregate annotations. While current compatibility predictions can prioritize therapeutic candidates, closing the gap to in vivo efficacy will likely require models that also incorporate in-gut persistence and amplification, and potentially community context, immune clearance, and tissue barriers..

### Surface architecture is the dominant axis of phage-host compatibility

Phage genus and K-locus-defined capsule type emerged as the strongest phage- and host-level determinants, respectively, placing the principal predictive signal at the cell surface. Phage genus represents lineage-associated traits including receptor-binding proteins, tail architecture, and other infection machinery. The substantial variance attributable to individual phage identity indicates that members of the same genus are not functionally interchangeable. On the host side, the capsule serves concurrently as a protective barrier to underlying structures and as an essential recognition substrate. Receptor-binding proteins and tailspikes can determine capsule specificity or shift host range when exchanged^29,31^, providing a clear mechanistic basis for this strong K-locus association.

This surface-first interpretation does not imply that capsule type specifies every terminal receptor. Some *Klebsiella* phages degrade the capsule or bypass by engaging O-antigen, FhuA, or other subcapsular structures^29,35,36^. Variation within capsule types may therefore reflect capsule expression, finer receptor specificity, secondary-receptor use, or post-adsorption compatibility. Similar patterns have been observed in other systematic interaction studies, including our companion studies in *E. coli*^61^ and *P. aeruginosa*^41^. By contrast, aggregate burdens of annotated defense systems, prophages, plasmid features, anti-defense genes, and antimicrobial-resistance determinants explained little of the binary susceptibility phenotype, although conditional effects may be obscured by a single clearance-based endpoint.

### Genome-wide features add resolution beyond curated descriptors

Although curated and learned representations capture overlapping biological mechanisms, they differ substantially in their predictive resolution, with both feature-rich models outperforming baseline predictors that rely solely on phage genus and capsule serotype. GenoPHI recovered this structure without curated host annotations, as its highest-ranked features map directly to capsule and LPS biosynthesis, canonical receptors, and defense- or prophage-associated modules. It also attributed a larger fraction of network variation to pair-specific signals than the curated baseline (32.4% versus 17.2%), suggesting that the raw pangenome features preserve fine-scale differences compressed or lost by aggregate annotations.

This additional genomic resolution was most valuable among closely related isolates with divergent susceptibility phenotypes. For ranking phages against a held-out isolate, however, GenoPHI improved AUROC only modestly and did not significantly improve AUPR. Its larger per-phage AUROC advantage indicates better discrimination among unseen hosts for phages represented during training, not generalization to novel phages. The weaker association between performance and phylogenetic novelty likely reflect the availability of close relatives for most cross-validation targets. GenoPHI therefore matches curated approaches on typical isolates, while providing necessary resolution for phenotypically atypical isolates. Its feature importance values, however, remain predictive associations rather than causal assignments.

### Cocktail testing defines the translational boundary of susceptibility prediction in *K. pneumoniae*

Empirical and genome-guided selection revealed a translational challenge between *in vitro* suppression and functional performance in the gut environment. Expert selection produced the most durable control in liquid culture, whereas only ML2 modestly reduced MKP103 colonization in mice. This divergence separates two core objectives in cocktail design, requiring researchers to concurrently combine phages that suppress growth and constrain escape, while ensuring that these entities remain viable, reach susceptible target cells, and amplify within the specific treatment environment. The association between bacterial reduction and fecal phage recovery implicates local exposure or replication as an important constraint in the gut, although the precise underlying mechanism remains unresolved. Modest efficacy *in vivo* cannot be attributed solely to inadequate host recognition, just as strong killing activity in well-mixed culture cannot be assumed to predict performance within a complex microbial community.

A conservative interpretation of this ML2 activity is warranted given that our cocktail selection approaches used different evidence streams, only four formulations were evaluated against a single strain, receptor assignments remained incomplete, and small animal groups alongside possible cage effects restricted generalization. Nevertheless, identifying an active cocktail for a bacterial isolate completely excluded from model training establishes the definitive feasibility of genome-guided prioritization when isolate-specific data are unavailable. These findings favor a hybrid clinical workflow that leverages genome-based ranking to narrow large pools of candidates, emphasizes receptor or resistance diversity to minimize shared pathways of mutational escape, and integrates targeted secondary screening of quantitative killing, regrowth kinetics, and amplification dynamics under treatment-relevant conditions.

Specific limitations bound the interpretation of these findings. The clearance-based solid-media spot assay primarily captures binary surface compatibility rather than quantitative or productive infection phenotypes, potentially underweighting downstream defense effects and limiting extrapolation to liquid culture or translational settings without further multi-environment validation. Statistical inference was also constrained by sparse representation and the necessary pooling of K-locus types for certain analyses, a reliance on genomic annotations rather than measured phenotypes, and the specific taxonomic composition of the current phage panel.

### Outlook

Future studies should expand host and phage diversity while incorporating real-time infection dynamics, resistance emergence kinetics, multi-environment stability, and site-specific replication parameters directly into model training. Implementing direct receptor prediction could effectively guide the assembly of cocktail combinations that engage independent structures, thereby limiting cross-resistance. Furthermore, evaluating these interactions within polymicrobial communities and biofilms would test therapeutic activity under realistic ecological constraints. In the longer term, this computational framework may guide the precise engineering of receptor-binding proteins to redirect viral tropism. Integrating interpretable genome-scale prediction with targeted phenotypic validation advances rapid phage prioritization for *K. pneumoniae* while defining the exact evidence needed to translate initial genetic compatibility into robust therapeutic formulations.

## Materials and Methods

### Bacterial Cultivation and Phage Acquisition

*Klebsiella pneumoniae* MKP103 was obtained from the Manoil Lab at the University of Washington, while the MRSN panel strains were sourced from the Walter Reed Army Institute of Research^51^. Culturing was performed at 37°C using LB agar (LB Lennox with 1.5% Bacto agar) or liquid LB (Lennox; 5 g/L NaCl, 5 g/L yeast extract, 10 g/L Tryptone).

Bacteriophages used in this research were isolated in-house or obtained from the KlebPhaCol collection. To determine phage titers, ten-fold serial dilutions were prepared in the SM buffer. Subsequently, 2 µL of each dilution was spot-plated onto 0.5% top agar lawns (5 g/L Bacto agar, 10 g/L NaCl, 10 g/L Tryptone) containing 5 mM MgSO₄ and 10 mM CaCl₂.

### Phage enrichment and amplification

2 mL of pooled environmental samples were combined with an equal volume of 2× LB supplemented with 10 mM CaCl₂ and MgSO₄, and inoculated with a 1:50 dilution of overnight *K. pneumoniae* cultures in tissue culture tubes and incubated at 37°C, with shaking at 200 rpm. The resulting enrichment was clarified by centrifugation (10,000 x g, 10 min) and filter-sterilized. For phage isolation, 10 ul of clarified enrichment was spotted on the lawn of individual *K. pneumoniae* MRSN strains and spread with sterile paper. Plaques of distinct morphologies were picked with a sterile P200 pipet tip, transferred to a microcentrifuge tube with 500 µl SM buffer and subjected to two additional rounds of spotting and streaking to obtain a clonal isolate with uniform plaque morphology. Where morphological heterogeneity persisted after three rounds, plaques of each distinct type were picked and purified separately to confirm that each represented a stable, distinct phage.

To amplify the phages, overnight *K. pneumoniae* cultures were diluted 1:50 into 30 mL of LB (with 5 mM CaCl₂ and 10 mM MgSO₄) and incubated at 37°C and 200 rpm for 30 minutes to achieve early exponential growth. The cultures were then inoculated with 100 µL of purified phage stock and co-incubated for 3 hours. Finally, the samples were harvested, clarified via centrifugation (10,000 × g for 10 min), and filter-sterilized, with successful amplification confirmed through subsequent titering. For mouse experiments, 30mL phage stocks were concentrated over a VivaSpin 15 Turbo Centrifugal Concentrator (VS15TR02) in batches until the final volume reached <1mL and resuspended in PBS.

### Phage Sequencing

Using a modified Wizard DNA purification kit (Promega), phage DNA was extracted from the amplified lysates as previously detailed. Neochromosome Inc. managed the NGS library preparation and Illumina sequencing for the phage genomes. The extracted DNA underwent RNase I treatment prior to library preparation. Phage DNA was then fragmented, and partial Illumina adapters were integrated through tagmentation using the Illumina TDE1 enzyme. After adding Nextera-compatible unique dual indices, sample libraries were amplified via PCR. The NGS libraries were then sequenced on an Illumina NextSeq2000 platform, generating paired-end 300bp reads with a target coverage depth of 10x for each sample.

### Phage Genome Assembly, Characterization and Comparative Analysis

Adapters were first trimmed from raw reads using cutadapt v4.7^62^ within trim_galore v0.6.10 (default parameters)^63^, and reads or regions of lower quality were identified and trimmed with bbduk v39.06 (qtrim=r, trimq=20, maq=20)^64–66^. The library was then normalized to 100x coverage with bbnorm v39.06 (target=100 min=2)^64–66^, before assembly with SPAdes v3.15.5 (--isolate -k 21,33,55,77,99,121)^67^. The resulting contigs of ≥2,000 bp were screened for phage genomes with geNomad v1.7.6 (--conservative, genomad database version 1.3)^68^, and their quality and completeness were evaluated with CheckV v1.0.3 (default parameters)^69^. For each library, a single phage genome estimated to be 100% complete was obtained, and selected for further characterization.

Phage genomes were annotated with Pharokka 1.6.1^70^. Specifically, coding sequences (CDS) were predicted with PHANOTATE 1.5.1^71^, tRNAs were predicted with tRNAscan-SE 2.0^72^, tmRNAs were predicted with Aragorn^73^ and CRISPRs were predicted with CRT^74^. Functional annotation was generated by matching each CDS to the PHROGs^75^, VFDB^76^ and CARD^77^ databases using MMseqs2^78^ and PyHMMER^79^. Contigs were matched to their closest hit in the INPHARED database^80^ using mash^81^. Annotations of coding genes were additionally curated with PHOLD v 0.2.0^82^.

Taxonomy was assigned to newly sequenced phages using vConTACT3 with reference database v.230^83^. Phage genome similarity was estimated using the proteomic equivalence quotient (PEQ) metric calculated by PhamClust^84^ for gene families inferred by PhaMMSeqs^85^. Briefly, the PEQ score for a pair of genomes is calculated as a product of the proportion of shared genes and average amino acid identity across all pairs of homologous proteins. The resulting PEQ similarity matrix of 84 phages was converted into a distance matrix in PHYLIP format and a distance-based phylogenetic tree was calculated by the FastME 2.0 online tool^86^. Complete *K.pneumoniae* public phage genomes from the vConTACT reference database v230 were generated with vConTACT3; most genus-level diversity documented among USA-reported were represented in our panel. Coverage gaps were confined to four low-frequency genera, each represented by only one or two assemblies in NCBI Virus.

### Bioinformatic analysis of *K. pneumoniae* Genomes

Genome assemblies for *K. pneumoniae* MRSN panel strains were retrieved from NCBI Datasets (BioProject accession PRJNA717739). Phage defense systems were identified using DefenseFinder 2.0.0^87^. Antibiotic resistance genes were identified using AMRFinder+ 3.11.11 with database version 2023-04-17.1^88^. Prophages were identified using geNomad 1.9.0^68^. Plasmid contigs and plasmid features, including replicon/incompatibility types, replication (rep), mobilization (mob), and conjugation genes, origin of transfer (oriT) sequences, circularity, and replicon distribution scores, were identified using Platon 1.6.0^89^. Capsule and O serotypes were predicted with Kaptive 2.0.7^90^. Core pangenome multiple nucleotide alignments were generated by PPanGGOLiN 2.0.3^91^ to calculate a *K. pneumoniae* phylogenetic tree via FastTree 2^92^.

### Phage host range determination

Phage-host interaction assays were performed by spotting 2 µL of each phage onto *K. pneumoniae* lawns using a semi-automated Rainin MicroPro 20 system. The 84 phages were tested against 101 strains, generating 8,484 interaction datapoints. Applied phage input ranged from 10^8^ to 10^11^ PFU/mL. Interactions were assessed based on the basis of host clearance. As mentioned in the results section, clearance can result from mechanical or enzymatic lysis, growth inhibition through toxicity or phage replication and productive lysis and in *K. pneumoniae* it can additionally arise from capsule depolymerisation, in which enzymatic degradation of the capsule produces halos and lawn clearing without a completed lytic cycle and from abortive infection, in which the cell dies without releasing progeny. These mechanisms cannot be reliably distinguished from single-titre spot assays. We nonetheless adopted a clearance-based definition of a positive interaction because the resulting phenotypes are consistent, strain-specific and directly relevant to therapeutic clearance of the pathogen, including in the capsule and biofilm contexts where depolymerase activity is itself beneficial. Interactions were therefore scored as 0 (no visible clearance), 1 (hazy/turbid clearance), or 2 (clear), averaged across two or more biological replicates, and classified as positive when the mean score reached ≥1.

### Statistical Analysis

#### Dataset and binarization

The phage-host interaction dataset comprised 101 *K. pneumoniae* strains tested against 84 phages, yielding an interaction matrix of 8,484 pairs. Because values 1 and 2 both represent interaction between bacteria-phage pairs with no meaningful biological distinction between interaction intensities in this dataset, all non-zero values were collapsed to 1, producing a binary classification target. The resulting class balance was 68.7% negative (5,828 pairs) and 31.3% positive (2,656 pairs). The pair-level dataset was constructed by melting the binary interaction matrix and merging phage metadata with selected strain-level features, producing one row per phage–host combination.

### Univariate phage-side analyses

Host range was defined as the number of strains interacting with a phage and was calculated by summing binary infection outcomes across the 101-strain panel. Two-sided Spearman rank correlations were used to evaluate univariate associations of host range with phage genome size, coding sequence count, and tRNA count. Genus-level differences in host range were assessed using a Kruskal–Wallis H test restricted to the nine genera represented by at least two phages (76 phages); eight singleton genera were excluded. Effect size was estimated as eta-squared, η² = (H − k + 1)/(N − k), where H is the Kruskal–Wallis statistic, k is the number of groups, and N is the number of observations included in the test. When the omnibus test was significant, pairwise differences between genera were evaluated using two-sided Dunn tests with Bonferroni adjustment.

### Univariate strain-side analyses

Per-strain interaction breadth or susceptibility was defined as the number of phages interacting with a strain and was calculated by summing binary infection outcomes across the 84-phage panel. Differences in susceptibility among the 54 K-serotypes were assessed using a Kruskal–Wallis H test, with η² calculated as described above. Because 31 serotypes were represented by one strain and 11 by two strains, the serotype analysis was treated as exploratory owing to sparse group sizes. Two-sided Spearman rank correlations were used to evaluate univariate associations of susceptibility with total prophage count; counts of prophages exceeding 75%, 90%, and 99% completeness; total gene count; plasmid length and coding capacity; counts of plasmid replicons, mobility genes, origins of transfer, and conjugation genes; plasmid-borne and chromosomal antimicrobial-resistance gene counts; and defense-system, anti-defense, and combined defense counts.

### Partial correlations

To assess whether each univariate association persisted after adjustment for selected covariates, Spearman partial correlations were computed. For continuous covariates, partial correlations were obtained with pingouin using the Spearman method; for categorical covariates (phage genus and K/capsule serotype), both the predictor and the response were first residualized by OLS regression on the categorical variable, and the Spearman correlation between the residuals was taken. On the phage side, the correlations of genome size, CDS count, and tRNA count with host range were each recomputed controlling for genus. On the strain side, prophage count was tested against susceptibility after adjustment for defense and antidefense counts; antidefense count was adjusted for prophage count; and defense count was adjusted for prophage and antidefense counts. A separate analysis tested defense count after adjustment for K/capsule serotype. Plasmid- and AMR-related features were additionally tested after adjustment for K-serotype, prophage count, defense count, antidefense count, and total gene count.

### Per-genus tests

To test whether genome-content associations with host range held within phage lineages, within-genus Spearman correlations between each genome feature (genome size, CDS count, and tRNA count) and host range were computed separately for each genus containing ≥4 phages (6 genera). Features that were invariant within a genus (e.g., tRNA count) were skipped, yielding 14 valid tests of 18 attempted. P-values were corrected for multiple testing using the Benjamini–Hochberg FDR procedure, applied both within each feature and pooled across all valid tests.

### Within- versus between-genus regressions

To distinguish associations arising from differences among genera from those arising among phages within the same genus, separate within–between linear regression models were fitted for genome size, CDS count, and tRNA count. Each genome feature was standardized across all 84 phages and decomposed into a between-genus component, defined by the genus mean, and a within-genus component, defined as each phage’s deviation from its genus mean. Host range was regressed simultaneously on the standardized between- and within-genus components. Standard errors were clustered by genus, and t-based inference used the 17 genus clusters (16 degrees of freedom). Regression coefficients were expressed as the change in host range per one-standard-deviation increase in the corresponding component

### Collinearity diagnostics

Multicollinearity among strain-level predictors was assessed using ordinary variance inflation factors calculated with variance_inflation_factor in statsmodels for the susceptibility-related predictor set and generalized variance inflation factors calculated with car::vif in R for the combined strain-profile predictor set. VIF and GVIF values < 5 and <2.24 respectively were taken to indicate moderate, interpretable collinearity.

### Type III ANOVA on susceptibility count

Strain susceptibility, defined as the number of infecting phages, was analyzed by Type III ANOVA using statsmodels. K- and O-serotype levels represented by fewer than three strains were lumped into an “Other” category. The model included the lumped K- and O-serotype factors together with prophage burden, plasmid length, plasmid replicon, mobility, origin-of-transfer and conjugation counts, plasmid- and chromosome-associated AMR counts, defense and antidefense counts, and total gene count. Partial η2 point estimates were calculated for each term, and overall model fit was summarized using R2, adjusted R2, and the model F-test.

### Crossed random-effects binomial GLMM

Binary interaction outcomes were analyzed at the individual strain–phage-pair level using crossed random-effects binomial generalized linear mixed models with a logit link, fitted with glmmTMB in R. The analysis included all 8,484 combinations of 101 strains and 84 phages. Prophage burden, plasmid length, plasmid replicon, mobility, origin-of-transfer and conjugation counts, plasmid- and chromosome-associated AMR counts, defense and antidefense counts, and total gene count were standardized and included as fixed effects. Full-resolution K-serotype, O-serotype, phage genus, and phage identity were represented by crossed random intercepts. A variance-decomposition model without a strain random intercept was used to partition latent-scale variation among the K-serotype, O-serotype, phage-genus, phage-identity, and logistic residual components; component intraclass correlations were calculated using π2/3 as the latent residual variance.

### Phylogeny-controlled PERMANOVA on susceptibility pattern

The pattern of strain susceptibility in the binarized matrix was analyzed by partial distance-based redundancy analysis (dbRDA) on Jaccard distances of per-strain infection profiles. To control for strain phylogenetic structure, the first 10 principal coordinates of neighbor matrices (PCNM eigenvectors) derived from the marker-gene phylogeny were entered as conditioning variables. The combined model was fitted with vegan::capscale and included lumped K- and O-serotype, plasmid- and chromosome-associated AMR counts, AMR phenotype, defense and antidefense counts, prophage burden, the number of prophages exceeding 90% completeness, and plasmid length, replicon, mobility, origin-of-transfer, and conjugation counts. Marginal permutation tests with 9,999 permutations were used to estimate each predictor’s unique contribution after adjustment for the remaining predictors and phylogenetic structure. Unique variance fractions were calculated as the marginal sum of squares divided by the total variance. Secondary adonis2 analyses tested each predictor separately while controlling for the first 10 PCNM axes. These secondary analyses did not control simultaneously for the other biological predictors. A separate combined adonis2 analysis used the full, non-lumped K- and O-serotype assignments to assess sensitivity to rare-serotype grouping.

### Phylogeny-controlled dbRDA on phage infection profiles

Phage-side variance in infection profiles was partitioned by distance-based redundancy analysis (dbRDA) on binary Jaccard distances between phages, using vegan::capscale. Two phages with no observed infections were excluded, leaving 82 phages shared among the interaction matrix, metadata, and corrected 84-tip PEQ tree. The PEQ-based phage phylogeny was decomposed into PCNM eigenvectors; the first 10 were entered as conditioning variables, and phage genus was the focal predictor. The phylogeny-adjusted contribution of the genus was quantified as constrained variance divided by total variance. Genus-only and phylogeny-only models were also fitted for comparison, and significance was assessed via 9,999 permutations.

### Phylogenetic signal

Phylogenetic signal for selected strain-level traits, including defense system count, antidefense count, prophage burden, susceptibility count, and seven plasmid-related features was quantified using Pagel’s λ on the marker-gene phylogeny, computed via phytools::phylosig (method=“lambda”, test=TRUE) in R. Likelihood-ratio tests against λ = 0 were used to assess significance.

### Two-way identity ANOVA

Binary interaction outcomes for all 8,484 combinations of 101 strains and 84 phages were analyzed using an additive two-way ordinary least-squares ANOVA with phage and strain identities as categorical fixed effects. Because the matrix was complete and balanced, the phage and strain main-effect Type II sums of squares were expressed as percentages of the corrected total variance. The phage-by-strain interaction was omitted because one observation per combination would saturate the model. The residual here therefore combines non-additive pair-specific interaction structure with assay/readout variation.

### Network analysis

We evaluated the bipartite phage–strain interaction network using modularity and nestedness. Modularity, which identifies groups with more within-group interactions than expected, was assessed using label propagation with bipartite modularity refinement (LP-BRIM)^93,94^, implemented in Python 3 using functions adapted from BiWeb^95^. Runs were refined to maximize Barber’s bipartite modularity Q^96^; 100 restarts were performed and the highest-Q partition was retained. ER72 and ER75 had no positive interactions and were excluded, leaving 101 strains, 82 phages, and 2,656 interactions. Realized modularity Q_r_ was the fraction of interactions within matching modules^97^. Nestedness was measured using strict NODF^98^, which tests whether phages with fewer interactions associated with subsets of the strains associated with phages having greater interaction breadth; combined, strain-wise, and phage-wise values were calculated from the complete 101 × 84 matrix, including ER72 and ER75.

Significance was assessed using 1,000 randomized matrices that preserved the interaction breadth of every strain and phage while rearranging their partners. For modularity, null matrices were generated using 5 × E row-pair trade attempts and five LP-BRIM restarts per replicate; the one-sided P-value was the proportion of null Q values at least as large as observed. For NODF, null matrices were generated using 5 × E pairwise swap attempts, and lower-than-expected nestedness was evaluated using the corrected lower-tail probability, (n_null≤observed_ + 1)/(N_null_ + 1). Standardized effect sizes were calculated as (observed − null mean)/null standard deviation.

Seeds were 42 for the observed modularity analysis, 123 for its null analysis, and 42 for the NODF null analysis. Module permeability was summarized for the fixed four-module partition by comparing the observed interaction density in each strain-module × phage-module block with that expected from strain and phage interaction breadths; enrichment was expressed as log2(observed/expected density).

### Jaccard distances and ordination

Jaccard distance matrices were computed from binary interaction scores using vegan::vegdist. Principal coordinate analyses (PCoA; k = 10) used stats::cmdscale. Phylogenetic and PEQ-based trees were converted to PCNM eigenvectors for use as conditioning variables in dbRDA and PERMANOVA models as described above.

## Machine learning modeling

### Models

We trained three deployable CatBoost classifiers on the binary interaction matrix under a 20-fold (10 folds × 2 reshufflings) strain-held-out cross-validation scheme, with 10% of strains entirely withheld from feature selection, training, and hyperparameter tuning in each fold:

- Genus + K-Serotype: a minimal-feature baseline using only phage genus (one-hot encoded) and host K-antigen capsule serotype (one-hot encoded). A variant additionally including O-antigen serotype was tested and found to be statistically indistinguishable from the K-only version (all paired Wilcoxon p ≥ 0.12); the K-only model is reported throughout.
- Curated baseline: adds strain-level defense system count, anti-defense system count, prophage burden, plasmid load, AMR phenotype, and genome-scale features (genome size, CDS count, gene count, virulence score), plus phage-level genome size, CDS count, and tRNA count.
- GenoPHI: replaces the curated bioinformatic features with pangenome-derived MMSeqs2 protein-cluster features and applies the model-based feature selection procedure described previously⁴⁵.

Within each fold, an ensemble of 50 CatBoost classifiers was trained on internal 80/20 splits of the training strains with early stopping on the internal held-out split. For each ensemble member, the best hyperparameters were chosen by an 8-point grid search (iterations ∈ {500, 1000}, learning_rate ∈ {0.05, 0.1}, depth ∈ {4, 6}) by MCC on the internal eval split, matching the per-run grid search used in GenoPHI’s feature-selection pipeline so that all three model classes received equivalent tuning budgets. All models used balanced class weights and Logloss as the training objective. Predictions on the held-out validation strains were summarized as the median confidence across ensemble members.

### Performance evaluation

Performance was evaluated under three aggregation views: per-fold (predictions pooled within each of the 20 folds), per-strain (predictions pooled across all folds in which each strain appeared, one AUROC/AUPR per strain), and per-phage (predictions pooled across all folds in which each phage appeared, one AUROC/AUPR per phage). For each view we computed AUROC, AUPR, and Matthews correlation coefficient (MCC). MCC and other threshold-dependent metrics were not used for per-phage comparisons because both deployable baselines collapsed to a single predicted class for the majority of phages under default thresholding (the curated baseline predicted no infection for ∼70% of phages), making per-phage MCC uninformative; per-phage comparisons are reported using AUROC and AUPR only. Pairwise model comparisons used paired Wilcoxon signed-rank tests within each aggregation view and metric.

### Difficulty-axis stratification

To localize where model performance differences are concentrated, per-strain AUROC, AUPR, and MCC were binned along two largely independent difficulty axes: (1) phylogenetic novelty, defined as patristic distance from the held-out strain to its nearest training-fold neighbor on a core-genome phylogeny; and (2) phenotypic divergence, defined as mean Jaccard distance between the held-out strain’s interaction profile (restricted to training-fold phages) and those of its five nearest phylogenetic neighbors in the training fold.

Both axis values were computed per (strain, iteration), then aggregated to the strain level as the median across iterations. Each strain was assigned to a tercile (low/mid/high) per axis using global cutpoints pooled across all models, and per-tercile performance was compared between models with paired Wilcoxon signed-rank tests. The phylogenetic novelty axis turned out to be largely uninformative because the maximum nearest-neighbor distance attainable on the strain tree (7.6 × 10⁻³) falls well below the median pairwise distance (1.0 × 10⁻²), so every held-out strain has a close training-set relative regardless of how random splits are drawn. We did not consider clade-aware splitting strategies, which would simulate discovery of a phylogenetically novel isolate rather than the routine clinical case of cross-validation targets.

### Cocktail design and benchmarking

For each deployable model, candidate 3- and 5-phage cocktails were selected per held-out strain using model confidence scores under receptor-diversity constraints derived by HDBSCAN clustering of the training-fold interaction matrix; the same clustering basis was used for all three deployable models to ensure model-to-model comparability. Cocktails were additionally compared against two non-ML baselines: (i) a greedy set-cover that iteratively selects the phage maximizing additional strain coverage on the training-fold interaction matrix; and (ii) a promiscuity baseline that hierarchically clusters phages on the training-fold interaction matrix and selects the most-infectious phage per cluster. Cocktail success was defined as ≥1 phage in the top-k recommendation that infects the held-out strain. Strains were stratified into quartiles by full-matrix susceptibility, and top-k success was compared between models within each quartile using McNemar’s test on paired strain-level outcomes.

### SHAP analysis

Shapley Additive exPlanations (SHAP) values were computed for the GenoPHI model using the TreeSHAP algorithm. The 25 strain-side feature clusters with the largest mean absolute SHAP value were annotated with gene functions using eggNOG-mapper to compare against known mediators of phage-host interaction in *K. pneumoniae*.

#### *In vivo* feasibility test using mice gut decolonization model

Germ-free Swiss-Webster mice were bred and housed within the University of Chicago Gnotobiotic Research Animal Facility (GRAF) in Trexler-style flexible firm isolators (Class Biologically Clean) within Ancare polycarbonate mouse cages (N10HT). Gnotobiotic mice were weaned at 21 days of age and were fed an autoclaved, plant-based mouse chow (LabDiet JL Rat and Mouse/Auto 6F 5K67). All mice were housed in cages containing autoclaved Teklad Pine Shavings cat# 7088) with a 12-hour light/dark cycle at a standard room temperature of 20-24 C. All mice were euthanized by CO₂ asphyxiation followed by cervical dislocation as a secondary measure. All experiments utilized both male and female mice. All experiments were performed in accordance with the Guide for the Care and Use of Laboratory Animals (8th ed.) and were approved by the Institutional Animal Care and Use Committee of the University of Chicago (Protocol 72610).

8-12 week old mice (control, ES2, ML1, and ML2) and 20 week old mice (ES1) were colonized with a defined microbial consortia (*Akkermansia muciniphila*, *Alistipes finegoldii*, *Anaerostipes caccae*, *Bacteroides thetaiotaomicron*, *Bacteroides uniformis*, *Bifidobacterium adolescentis*, *Escherichia coli* Nissle, *Limosilactobacillus reuteri*, *Parabacteroides distasonis*, *Phocaeicola vulgatus*, *Roseburia inulinivorans*, *Ruminoccocus gnavus*, and *Subdoligranulum variable*) on days 1 and 4. On day 9, mice were gavaged with 100 µL *Klebsiella pneumoniae* MKP103 at 5 x 10^8^ CFU/mL in PBS and monitored by fecal collections. Cell titers were determined by resuspension of fecal pellets in 1 mL PBS and shaken for 2 minutes at 2,000 rpm in a PowerLyzer 24 homogenizer (Qiagen), followed by a quick 30 s spin at 300 g. 10-fold serial dilutions were spotted on LB carbenicillin plates at room temperature and enumerated after overnight incubation. On days 14 and 15, each mouse was gavaged with 100 µl freshly made 5% sodium bicarbonate followed by 100 µl cocktail or PBS. Phage titer was determined by serial dilution of the homogenized fecal slurry supernatant after a 10-minute spin at 10,000 g. ES1 and ES2 were enumerated on MKP103 and ER649, ML1 was counted on ER649 and 164125A, and ML2 on MKP103, ER649, and 164135A, with the higher titer being reported.

## Data Availability

Assembled genome sequences for all phages characterized in this study have been deposited in NCBI GenBank under accession numbers (being submitted). Matrix data and curated feature tables are listed in Supplementary Datasets (Figshare - 10.6084/m9.figshare.33235941).

## Code Availability

The GenoPHI software package used for phage-host interaction prediction is available at https://github.com/Noonanav/GenoPHI.

## Author contributions

S., Conceptualization, Methodology, Investigation, Validation, Formal Analysis, Data Curation, Visualization, Writing - Original Draft, Review & Editing

J C. N., Conceptualization, Methodology, Investigation, Validation, Formal Analysis, Data Curation, Visualization, Writing - Original Draft, Review & Editing

R., Methodology, Investigation, Validation, Formal Analysis, Visualization, Data Curation, Writing - Original Draft, Review & Editing

A., Investigation, Validation D.P., Investigation, Validation

M., Formal Analysis, Data Curation, Visualization

K. V., Investigation, Validation, Visualization

S., Investigation, Validation

O., Investigation, Validation

C., Investigation, Validation

M., Investigation, Validation

B., Investigation, Validation

K., Investigation, Software, Formal Analysis, Data Curation, Visualization, Writing - Original Draft, Review & Editing

M. D., Investigation, Writing - Review & Editing

R., Investigation, Software, Formal Analysis, Data Curation, Visualization, Writing - Original Draft, Review & Editing

M., Methodology, Investigation, Validation, Formal Analysis, Visualization, Writing - Original Draft, Review & Editing, Funding Acquisition

A.P. A., Conceptualization, Methodology, Writing - Review & Editing, Supervision, Funding Acquisition

K. M., Conceptualization, Methodology, Writing - Review & Editing, Supervision, Funding Acquisition, Project Administration

## Materials & Correspondence

All other data supporting the findings of this study are available from the corresponding author Vivek K. Mutalik or Adam P. Arkin upon reasonable request.

## Acknowledgments

We thank members of the entire BRaVE Phage Foundry team, the Arkin lab and the Mutalik lab for discussions.

