## Supplemental Information for "A comprehensive phage-bacteria interaction atlas links phage lineage and capsule serotype to genome-guided machine learning prediction in *Klebsiella pneumoniae*"

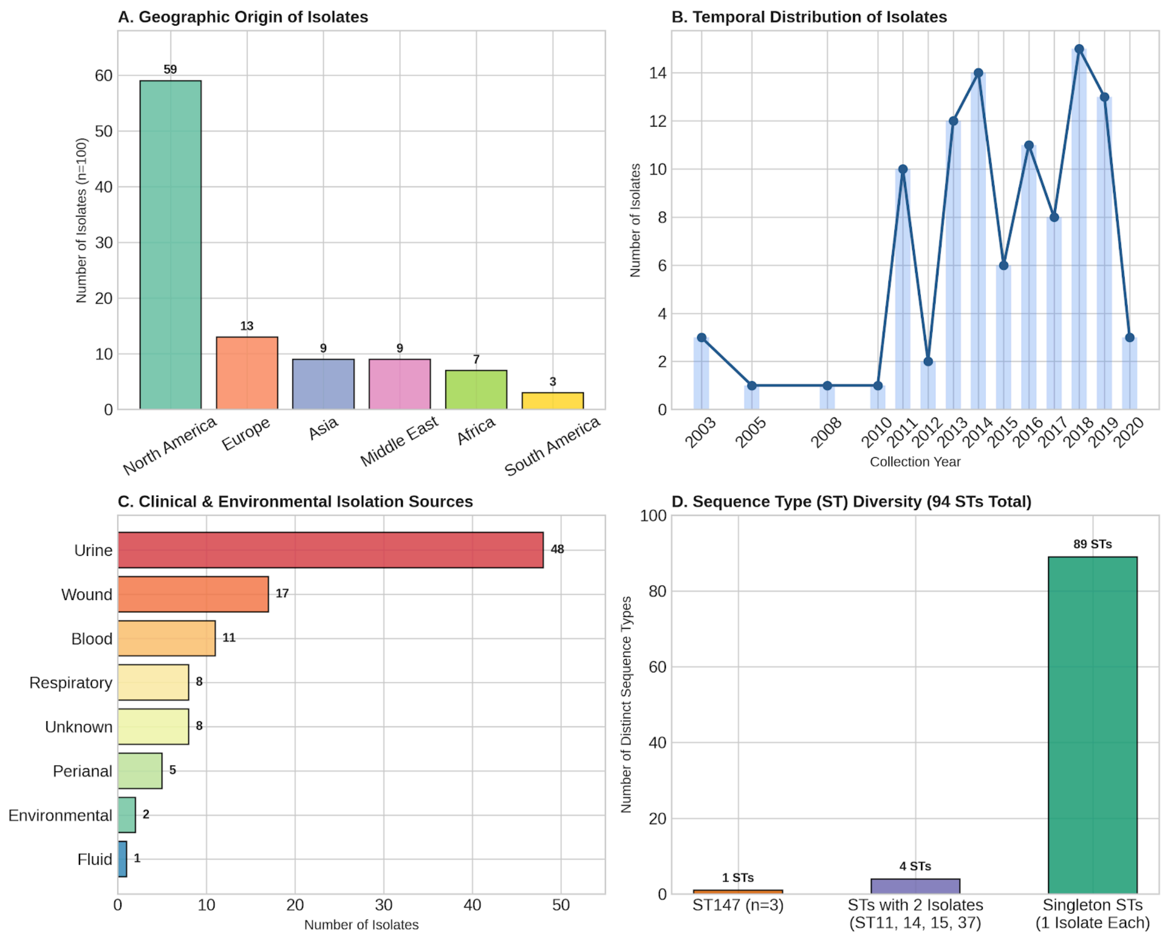


**Figure S1: Demographic, temporal, clinical, and sequence diversity of the *Klebsiella pneumoniae* MRSN panel (N = 100).** (A) Geographic distribution of panel isolates across six global regions. (B) Annual collection timeline of isolates spanning 2003 to 2020. (C) Frequency breakdown of isolates by clinical anatomic site and surveillance source. (D) High genomic sequence type (ST) diversity across the panel, comprising 94 unique MLST sequence types.

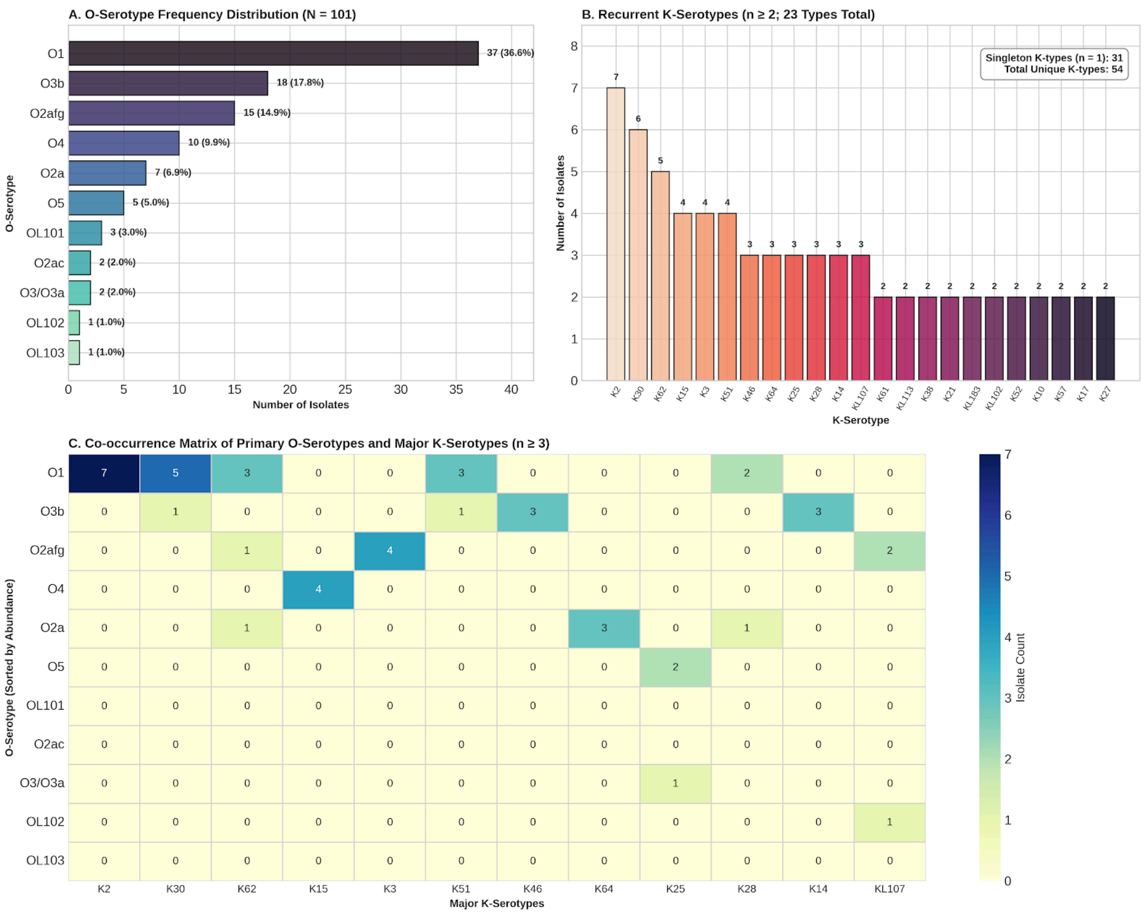


**Figure S2: Distribution and co-occurrence of K- and O-serotypes across the panel.** (A) Frequency distribution of the 11 identified O-lipopolysaccharide serotypes. Percentages denote the proportion of total isolates. (B) Frequency distribution of recurrent capsular K-serotypes (n>=2; 23 unique types). Inset indicates the total count of unique K-serotypes (n = 54) and singletons (n = 31). (C) Co-occurrence matrix showing the intersection between primary O-serotype backbones and major K-serotypes (n>=3). Values inside heatmap cells indicate exact isolate counts, highlighting specific K- and O-antigen pairings.

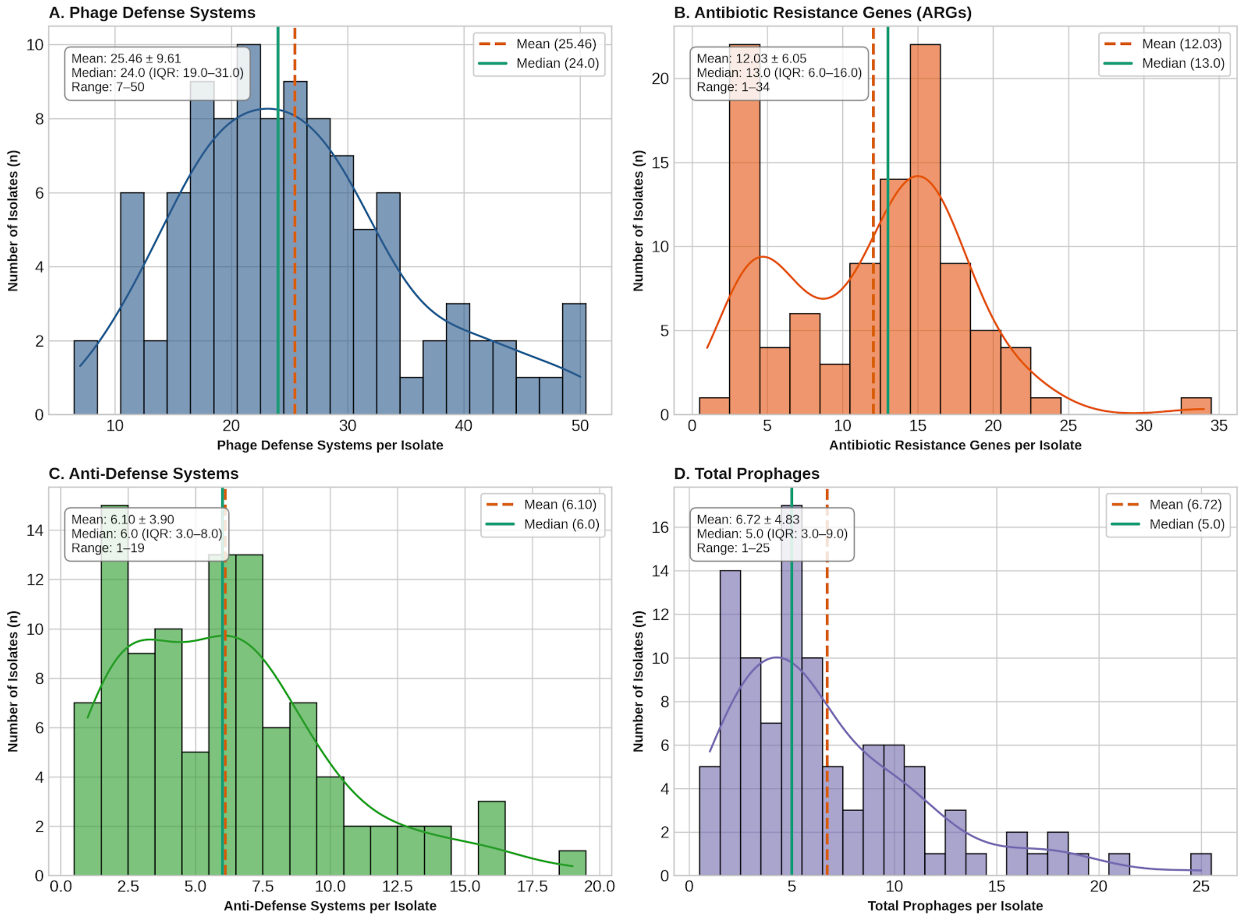


**Figure S3: Frequency distributions and central tendencies of phage defense systems, antibiotic resistance genes, anti-defense systems, and prophages across the *K. pneumoniae* panel (N = 101).** (A) Distribution of phage defense systems per genome (mean $= 25.46; median = 24). (B) Distribution of total antibiotic resistance genes (ARGs) per genome (mean = 12; median = 13). Peaks correspond to wild-type/susceptible strains (1–5 ARGs) and multidrug-resistant lineages (12–18 ARGs). (C) Distribution of anti-defense systems per genome (mean = 6.10; median = 6.0, IQR: 3.0–8.0). (D) Right-skewed distribution of total integrated prophages per genome (mean = 6.72; median = 5). In all panels, vertical dashed orange lines indicate the arithmetic mean and solid teal lines denote the sample median. Curves represent kernel density estimations.

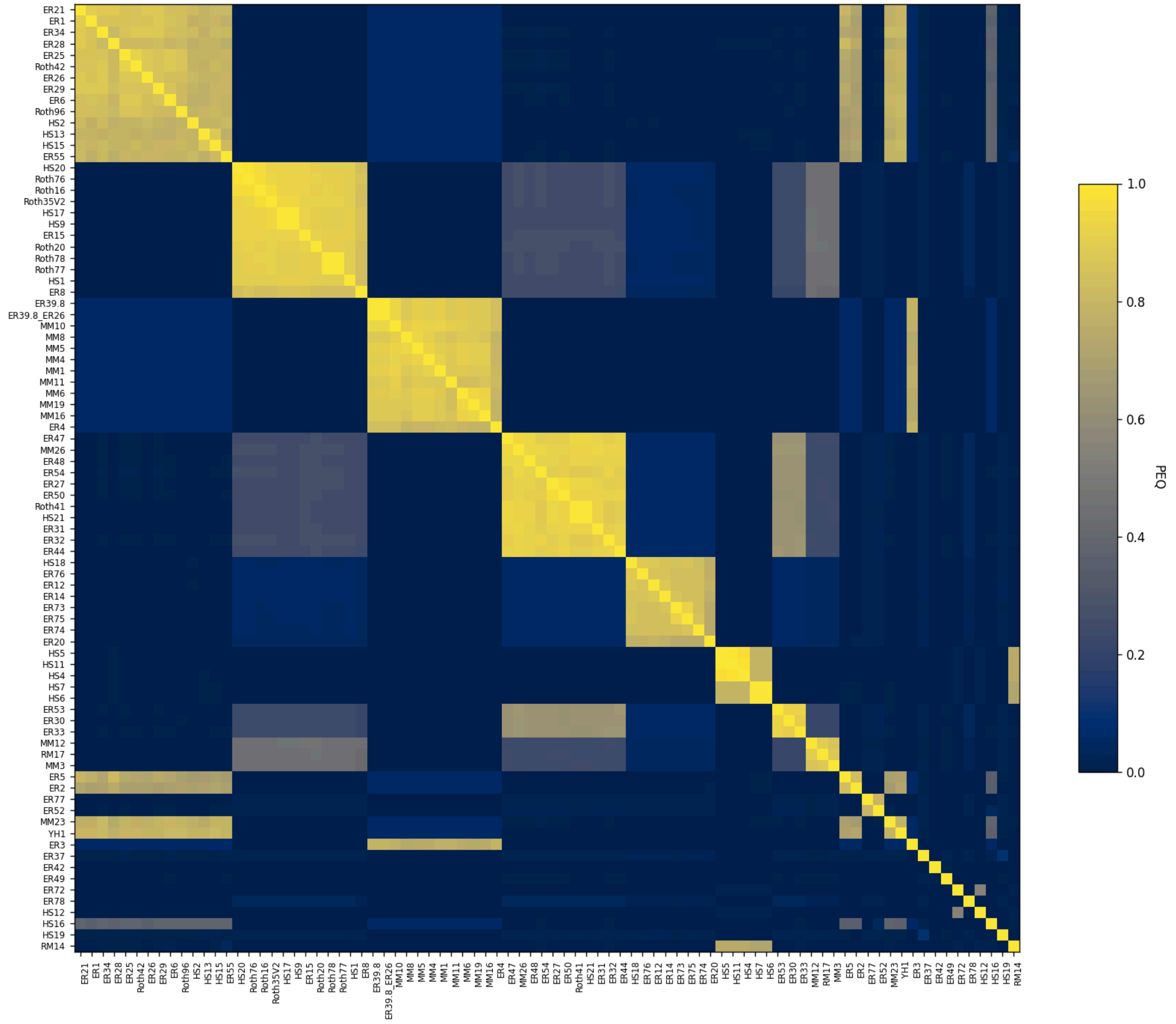


**Figure S4: PhamClust comparison of *K. pneumoniae* phages.** All *K. pneumoniae* phages (n=84) were sequenced and analyzed via PhaMMseqs and PhamClust. The heatmap shows the proteomic equivalent quotient (PEQ) of these phages. Phage names are listed along the X- and Y-axes.

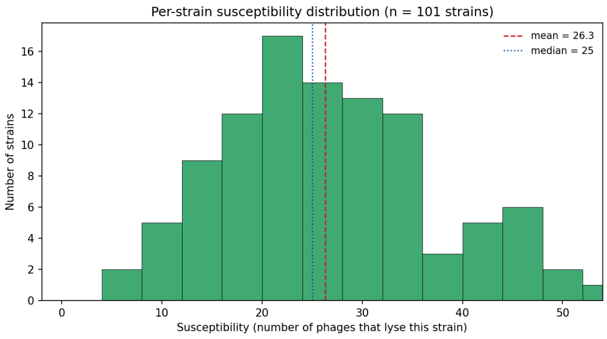


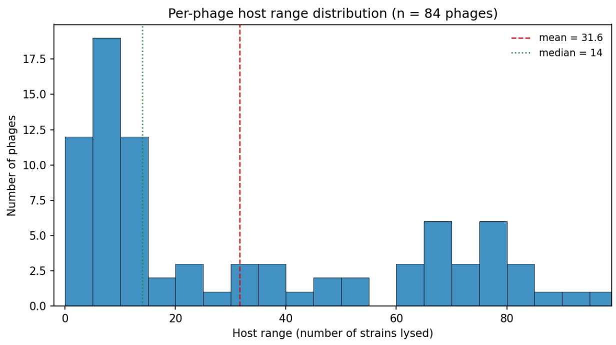

**Figure S5: Distribution of strain- and phage-level interaction breadth.** A) Per-strain interaction breadth or susceptibility among 101 *K. pneumoniae* strains, defined as the number of phages in the 84-phage panel that produced a positive interaction with each strain. Interaction breadth ranged from 6 to 52 phages (mean, 26.3; median, 25). Every strain interacted positively with at least six phages, although the number of positive interactions varied substantially among strains. B) Per-phage interaction breadth or host range among 84 phages, defined as the number of strains in the 101-strain panel that produced a positive interaction with each phage. Interaction breadth ranged from 0 to 97 strains (mean, 31.6; median, 14). The mean exceeding twice the median indicates that most phages interacted with relatively few strains, whereas a small subset exhibited broad interaction profiles across the strain panel. Two phages produced no positive interactions under the assay conditions. Red dashed lines indicate means, and dotted lines indicate medians.

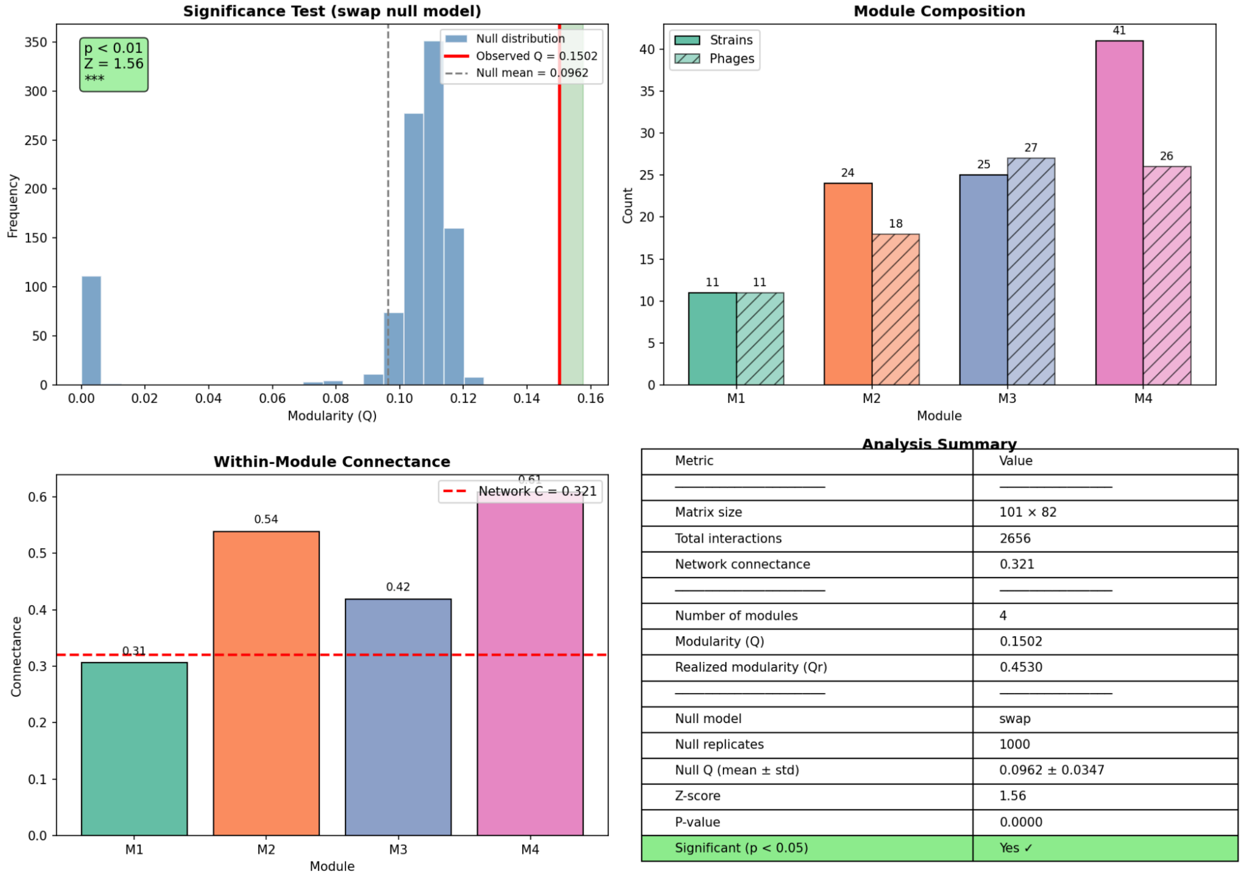


| Metric | Value |
|---|---|
| ———————— | ———————— |
| Matrix size | 101 × 82 |
| Total interactions | 2656 |
| Network connectance | 0.321 |
| ———————— | ———————— |
| Number of modules | 4 |
| Modularity (Q) | 0.1502 |
| Realized modularity (Qr) | 0.4530 |
| ———————— | ———————— |
| Null model | swap |
| Null replicates | 1000 |
| Null Q (mean ± std) | 0.0962 ± 0.0347 |
| Z-score | 1.56 |
| P-value | 0.0000 |
| Significant (p < 0.05) | Yes ✓ |

**Figure S6: Modularity analysis of the phage-bacteria interaction network.** Statistical significance of network modularity calculated using 1,000 swap null model iterations (A) alongside strain and phage composition (B) and within-module connectance (C) across four identified modules (M1–M4). Complete network topology and null hypothesis parameters are summarized in (D).

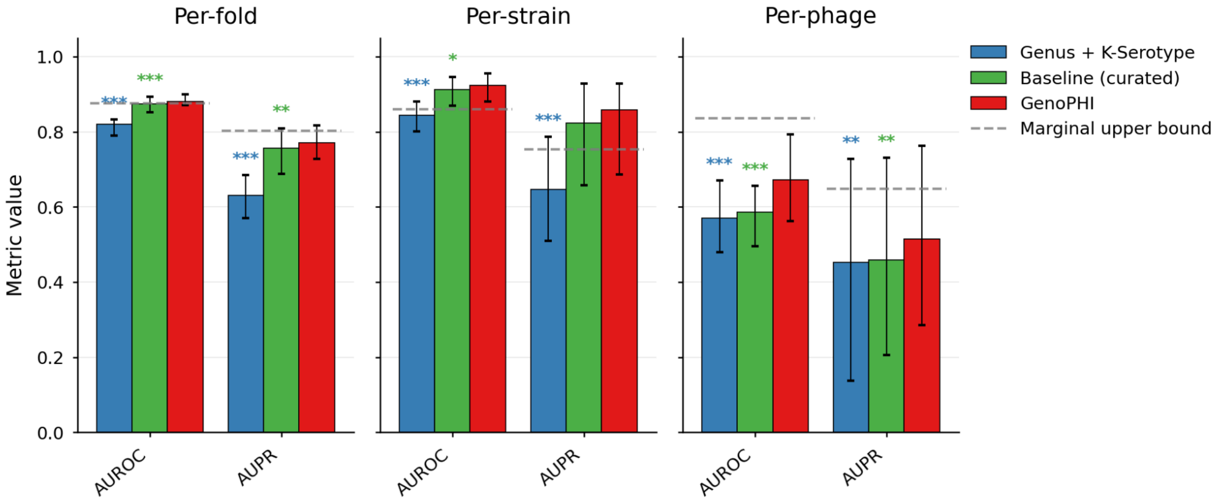


Supplementary Figure S7. Strain-versus-phage performance asymmetry. Per-entity performance for the three deployable models (Genus + K-Serotype, blue; curated baseline, green; GenoPHI, red) under 20-fold strain-held-out cross-validation, computed at three grains: pooled across all held-out predictions (Overall), per strain (AUROC/AUPR computed within each held-out strain over all phages, then summarized across strains), and per phage (computed within each phage over all strains). Bars show the median; error bars span the interquartile range. Asterisks denote Holm-corrected paired Wilcoxon significance versus GenoPHI (* $p < 0.05$, ** $p < 0.01$, *** $p < 0.001$). The two feature-rich models are close on per-strain ranking but diverge on per-phage ranking, where GenoPHI leads; both clearly exceed Genus + K-Serotype at every grain.

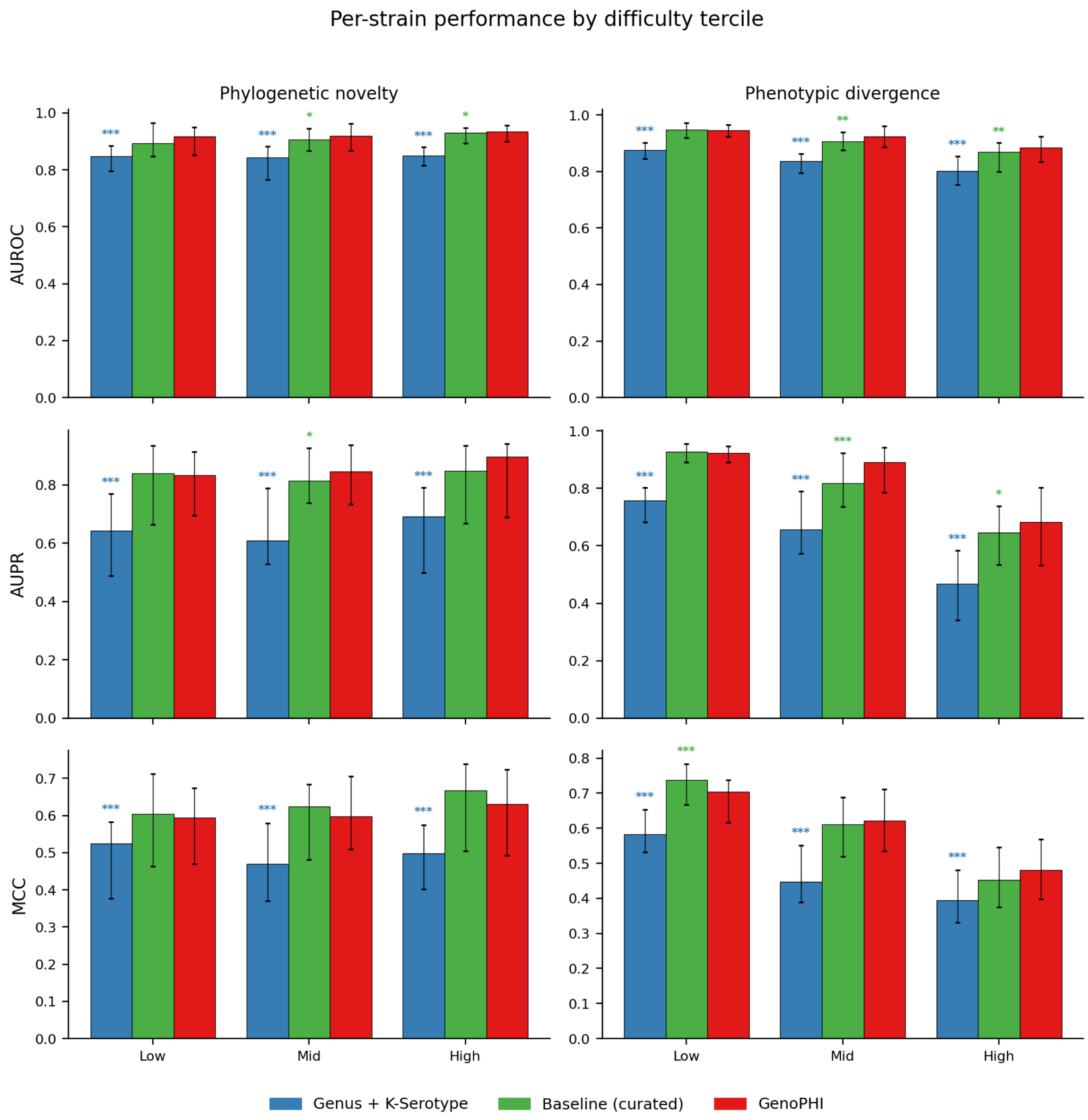


Supplementary Figure S8. Per-strain performance stratified by difficulty tercile. Per-strain held-out AUROC, AUPR, and MCC (rows) binned into terciles of two difficulty axes (columns): phylogenetic novelty (patristic distance from each held-out strain to its nearest training-set neighbor) and phenotypic divergence (mean Jaccard distance between a held-out strain's interaction profile and those of its five nearest phylogenetic neighbors in training). Terciles are defined on the pooled per-strain distribution of each axis. Bars show the median across strains for each model (Genus + K-Serotype, blue; curated baseline, green; GenoPHI, red); error bars span the interquartile range. Asterisks denote Holm-corrected paired Wilcoxon significance versus GenoPHI within each tercile (* $p < 0.05$, ** $p < 0.01$, *** $p < 0.001$). The GenoPHI-versus-baseline gap runs in the same direction along both axes but is weaker and less orderly along phylogenetic novelty, consistent with the compressed novelty range of this panel (maximum nearest-neighbor distance $7.6\times10^{-3}$ versus median pairwise $1.0\times10^{-2}$). The divergence-AUROC panel corresponds to Fig. 6C.

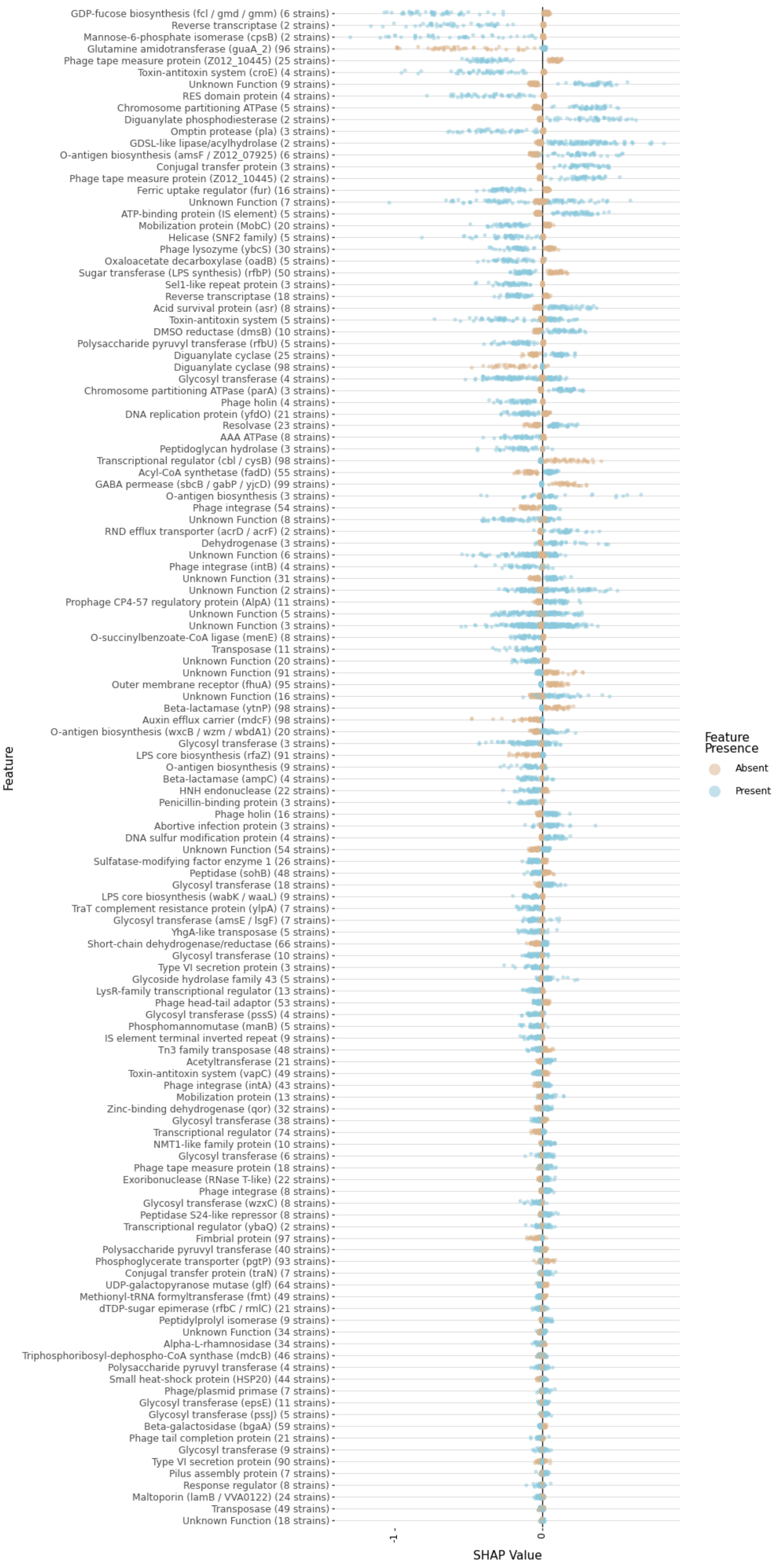
GDP-fucose biosynthesis (fcl / gmd / gmm) (6 strains)
Reverse transcriptase (2 strains)
Mannose-6-phosphate isomerase (cpsB) (2 strains)
Glutamine amidotransferase (guaA_2) (96 strains)
Phage tape measure protein (Z012_10445) (25 strains)
Toxin-antitoxin system (croE) (4 strains)
Unknown Function (9 strains)
RES domain protein (4 strains)
Chromosome partitioning ATPase (5 strains)
Diguanylate phosphodiesterase (2 strains)
Omptin protease (pla) (3 strains)
GDSL-like lipase/acylhydrolase (2 strains)
O-antigen biosynthesis (amsF / Z012_07925) (6 strains)
Conjugal transfer protein (3 strains)
Phage tape measure protein (Z012_10445) (2 strains)
Ferric uptake regulator (fur) (16 strains)
Unknown Function (7 strains)
ATP-binding protein (IS element) (5 strains)
Mobilization protein (MobC) (20 strains)
Helicase (SNF2 family) (5 strains)
Phage lysozyme (ybcS) (30 strains)
Oxaloacetate decarboxylase (oadB) (5 strains)
Sugar transferase (LPS synthesis) (rfbP) (50 strains)
Sel1-like repeat protein (3 strains)
Reverse transcriptase (18 strains)
Acid survival protein (asr) (8 strains)
Toxin-antitoxin system (5 strains)
DMSO reductase (dmsB) (10 strains)
Polysaccharide pyruvyl transferase (rfbU) (5 strains)
Diguanylate cyclase (25 strains)
Diguanylate cyclase (98 strains)
Glycosyl transferase (4 strains)
Chromosome partitioning ATPase (parA) (3 strains)
Phage holin (4 strains)
DNA replication protein (yfdO) (21 strains)
Resolvase (23 strains)
AAA ATPase (8 strains)
Peptidoglycan hydrolase (3 strains)
Transcriptional regulator (cbl / cysB) (98 strains)
Acyl-CoA synthetase (fadD) (55 strains)
GABA permease (sbcB / gabP / yjcD) (99 strains)
O-antigen biosynthesis (3 strains)
Phage integrase (54 strains)
Unknown Function (8 strains)
RND efflux transporter (acrD / acrF) (2 strains)
Dehydrogenase (3 strains)
Unknown Function (6 strains)
Phage integrase (intB) (4 strains)
Unknown Function (31 strains)
Unknown Function (2 strains)
Prophage CP4-57 regulatory protein (AlpA) (11 strains)
Unknown Function (5 strains)
Unknown Function (3 strains)
O-succinylbenzoate-CoA ligase (menE) (8 strains)
Transposase (11 strains)
Unknown Function (20 strains)
Unknown Function (91 strains)
Outer membrane receptor (fhuA) (95 strains)
Unknown Function (16 strains)
Beta-lactamase (ytnP) (98 strains)
Auxin efflux carrier (mdcF) (98 strains)
O-antigen biosynthesis (wxcB / wzm / wbdA1) (20 strains)
Glycosyl transferase (3 strains)
LPS core biosynthesis (rfaZ) (91 strains)
O-antigen biosynthesis (9 strains)
Beta-lactamase (ampC) (4 strains)
HNH endonuclease (22 strains)
Penicillin-binding protein (3 strains)
Phage holin (16 strains)
Abortive infection protein (3 strains)
DNA sulfur modification protein (4 strains)
Unknown Function (54 strains)
Sulfatase-modifying factor enzyme 1 (26 strains)
Peptidase (sohB) (48 strains)
Glycosyl transferase (18 strains)
LPS core biosynthesis (wabK / waaL) (9 strains)
TraT complement resistance protein (ylpA) (7 strains)
Glycosyl transferase (amsE / lsgF) (7 strains)
YhgA-like transposase (5 strains)
Short-chain dehydrogenase/reductase (66 strains)
Glycosyl transferase (10 strains)
Type VI secretion protein (3 strains)
Glycoside hydrolase family 43 (5 strains)
LysR-family transcriptional regulator (13 strains)
Phage head-tail adaptor (53 strains)
Glycosyl transferase (pssS) (4 strains)
Phosphomannomutase (manB) (5 strains)
IS element terminal inverted repeat (9 strains)
Tn3 family transposase (48 strains)
Acetyltransferase (21 strains)
Toxin-antitoxin system (vapC) (49 strains)
Phage integrase (intA) (43 strains)
Mobilization protein (13 strains)
Zinc-binding dehydrogenase (qor) (32 strains)
Glycosyl transferase (38 strains)
Transcriptional regulator (74 strains)
NMT1-like family protein (10 strains)
Glycosyl transferase (6 strains)
Phage tape measure protein (18 strains)
Exoribonuclease (RNase T-like) (22 strains)
Phage integrase (8 strains)
Glycosyl transferase (wzxC) (8 strains)
Peptidase S24-like repressor (8 strains)
Transcriptional regulator (ybaQ) (2 strains)
Fimbrial protein (97 strains)
Polysaccharide pyruvyl transferase (40 strains)
Phosphoglycerate transporter (pgtP) (93 strains)
Conjugal transfer protein (traN) (7 strains)
UDP-galactopyranose mutase (glf) (64 strains)
Methionyl-tRNA formyltransferase (fmt) (49 strains)
dTDP-sugar epimerase (rfbC / rmlC) (21 strains)
Peptidylprolyl isomerase (9 strains)
Unknown Function (34 strains)
Alpha-L-rhamnosidase (34 strains)
Triphosphoribosyl-dephospho-CoA synthase (mdcB) (46 strains)
Polysaccharide pyruvyl transferase (4 strains)
Small heat-shock protein (HSP20) (44 strains)
Phage/plasmid primase (7 strains)
Glycosyl transferase (epsE) (11 strains)
Glycosyl transferase (pssJ) (5 strains)
Beta-galactosidase (bgaA) (59 strains)
Phage tail completion protein (21 strains)
Glycosyl transferase (9 strains)
Type VI secretion protein (90 strains)
Pilus assembly protein (7 strains)
Response regulator (8 strains)
Maltoporin (lamB / VVA0122) (24 strains)
Transposase (49 strains)
Unknown Function (18 strains)
Feature
-1
0
SHAP Value
Feature Presence
Absent
Present

Supplementary Figure S9. Full GenoPHI SHAP feature set. SHAP beeswarm of all strain-side pangenome feature clusters retained by GenoPHI's model-based feature selection, extending the top-25 shown in Fig. 6E, annotated with eggNOG-mapper gene-function calls. Each point is one modeling run's median SHAP value for that cluster (50 runs), shown separately for its present (blue) and absent (orange) states; positive SHAP values increase predicted infection probability, negative values decrease it. Strain counts in parentheses give the number of strains in which each cluster is present. Beyond the top-ranked capsule and lipopolysaccharide biosynthesis clusters, the selected set includes canonical outer-membrane phage receptors, including the ferrichrome receptor fhuA and the maltoporin lamB.

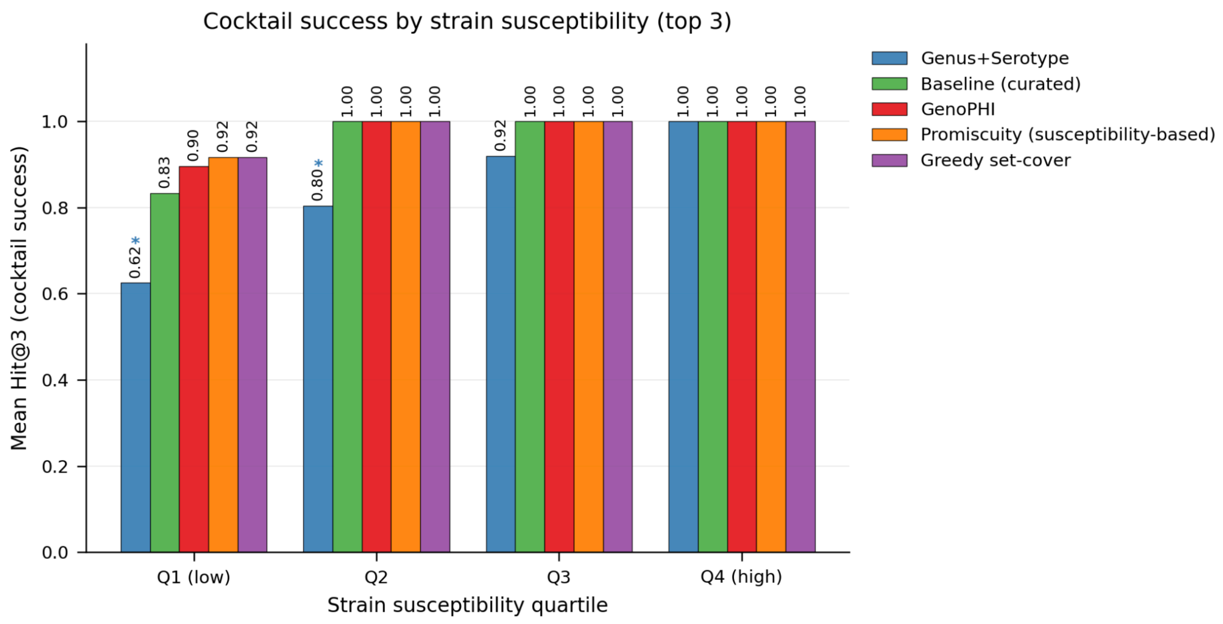


Figure S10: We benchmarked five cocktail-selection strategies under the same 20-fold cross-validation used for model evaluation (Fig. 7). Ranking phages by each model's confidence, we built 3- and 5-phage cocktails per held-out strain subject to receptor-diversity constraints derived by HDBSCAN clustering of the training-fold interaction matrix, then scored each model by whether its top-k recommendation contained at least one strain-infecting phage. Two non-ML procedures served as comparators: a greedy set-cover tuned to maximize strain coverage in the training fold, and a published promiscuity selector that clusters the training-fold interaction matrix hierarchically and orders phages by their within-cluster breadth in that fold. In the bottom susceptibility quartile (Q1), where correctly ranking the few infecting phages is decisive, the four rational strategies performed comparably: GenoPHI reached 89.6% top-3 success, statistically indistinguishable from greedy set-cover (91.7%), the promiscuity selector (91.7%), and the curated baseline (83.3%; all Benjamini-Hochberg-corrected McNemar $p \geq 0.99$), while Genus + K-Serotype trailed at 62.5% (corrected $p = 0.026$). At top-5 within Q1 the same ordering held, with GenoPHI at 91.7%, greedy set-cover highest at 97.9%, and Genus + K-Serotype lowest at 70.8% (differences no longer significant after correction). From Q2 through

Q4 every deployable approach other than Genus + K-Serotype reached 100% top-3 success, so cocktail design collapses to a coverage problem; Genus + K-Serotype remained behind in Q2 (80.4%, corrected $p = 0.043$) and converged only by Q4. This pattern is consistent with the modular network architecture established earlier: the low-susceptibility (Q1) strains have access to the narrowest range of compatible phages, so the cost of picking from the wrong module is highest and per-strain targeting matters most. Because the susceptibility profile of an incoming clinical isolate cannot be known in advance, a genome-based model that performs on par with coverage-optimized non-ML selectors on the hardest strains, while requiring no prior interaction data, is a practical option to advance toward clinical use.
